# Chemogenetic Manipulations of Ventral Tegmental Area Dopamine Neurons Reveal Multifaceted Roles in Cocaine Abuse

**DOI:** 10.1101/246595

**Authors:** Stephen V Mahler, Zachary D Brodnik, Brittney M Cox, William C Buchta, Brandon S Bentzley, Zackary A Cope, Edwin C Lin, Matthew D Riedy, Michael D Scofield, Justin Messinger, Arthur C Riegel, Rodrigo A España, Gary Aston-Jones

## Abstract

Ventral tegmental area (VTA) dopamine (DA) neurons perform diverse functions in motivation and cognition, but their precise roles in addiction-related behaviors are still debated. Here, we targeted VTA DA neurons for bidirectional chemogenetic modulation during specific tests of cocaine reinforcement, demand, and relapse-related behaviors, querying the roles of DA neuron inhibitory and excitatory G-protein signaling in these processes. Designer receptor stimulation of Gq-, but not Gs-signaling in DA neurons enhanced cocaine seeking via functionally distinct projections to forebrain limbic regions. In contrast, engaging inhibitory Gi/o signaling in DA neurons blunted cocaine’s reinforcing and priming effects, reduced stress-potentiated reinstatement, and altered cue-induced cocaine seeking strategy, but not the motivational impact of cocaine cues per se. Results demonstrate that DA neurons play several distinct roles in cocaine seeking, depending on behavioral context, G-protein signaling, and DA neuron efferent target, highlighting their multifaceted roles in addiction.

**Significance Statement:** G-protein coupled receptors are crucial modulators of VTA dopamine neuron activity, but how metabotropic signaling impacts dopamine’s complex roles in reward and addiction is poorly understood. Here, we bidirectionally modulate dopamine neuron G-protein signaling with DREADDs during a variety of cocaine seeking behaviors, revealing nuanced, pathway-specific roles in cocaine reward, effortful seeking, and relapse-like behaviors. Gq- and Gs-stimulation activated dopamine neurons, but only Gq stimulation robustly enhanced cocaine seeking. Gi/o inhibitory signaling altered the response strategy employed during cued reinstatement, and reduced some, but not all types of cocaine seeking. Results show that VTA dopamine neurons modulate numerous distinct aspects of cocaine addiction- and relapse-related behaviors, and indicate potential new approaches for intervening in these processes to treat addiction.

## Introduction

Ventral tegmental area (VTA) is a crucial node in mesolimbic circuits of reward, and the 50-70% of neurons there that express dopamine (DA)(Dobi et al., 2010) play diverse, and long-debated roles in natural and drug reward-related behaviors. The activity of these neurons is extensively regulated by G-protein coupled receptors (GPCRs), and several GPCRs are currently under investigation as therapeutic targets for the treatment of cocaine use disorders. Nevertheless, the specific roles for DA neurons in cocaine reinforcement, effortful seeking, or reinstatement, and how GPCR signaling modulates these, remain unclear.

Studies using pharmacological manipulation of DA receptor signaling, lesions of DAergic regions and their forebrain targets, or measurement of DA neuron firing or nucleus accumbens (NAc) DA release show clear roles for DA circuits in reward salience, learning, valuation, and seeking(Berridge and Robinson, 1998; Everitt et al., 2008; Bromberg-Martin et al., 2010; Luscher and Malenka, 2011; Lammel et al., 2014a; Keiflin and Janak, 2015; Hamid et al., 2016; Schultz, 2016). VTA DA neuron-specific optogenetic stimulation experiments confirm reinforcing and conditioned motivational roles for DA neurons; conversely, optogenetic inhibition of VTA DA neurons is behaviorally avoided(Tsai et al., 2009; Adamantidis et al., 2011; Witten et al., 2011; Steinberg et al., 2013; Berrios et al., 2016). Specific dissections of VTA DA neuron behavioral functions have also revealed marked functional heterogeneity, relating in part to efferent projection target and co-transmitter activity(Floresco and Magyar, 2006; Chaudhury et al., 2013; Lammel et al., 2014b; Trudeau et al., 2014; Barker et al., 2016; Tritsch et al., 2016). However, the roles for DA neurons in cocaine self-administration or relapse-like behaviors, and how GPCR signaling in these neurons affects these behavioral functions, are poorly understood.

Here, we used a chemogenetic approach to probe the roles of VTA DA neurons, and their main forebrain targets, in cocaine intake and relapse. Taking advantage of the transient, reversible modulation offered by Designer Receptors Exclusively Activated by Designer Drugs (DREADDs) to repeatedly and selectively manipulate DA neurons and their efferents during cocaine seeking behaviors, we demonstrate that engaging Gi- and Gq- (but not Gs) signaling in VTA DA neurons causes nonsymmetrical, task-dependent changes in drug seeking and cognition, depending on whether behavior was motivated by cocaine’s reinforcing or priming effects, cocaine-paired cues, or stress-like states.

## Materials and Methods

### Subjects

Male, hemizygous TH::Cre transgenic rats (n=130) and WT littermates (n=33) were bred at MUSC, UCI, or Drexel Universities, from founders provided by K. Deisseroth (mated to wild type partners). They were housed in tub cages under reverse 12:12 light cycle, with free access to food and water at all times (initial surgical weight 250-400g). All procedures were approved by the MUSC, UCI, or Drexel institutional animal care and use committees.

### Viral Constructs

Immediately after catheter implantation, 1µl/hemisphere of an AAV2 vector containing a double-floxed, inverted open reading frame (DIO) sequence for the mCherry-tagged hM4Di (Gi-coupled), hM3Dq (Gq-coupled), or rM3Ds (Gs-coupled) DREADD, or mCherry control (*hSyn-DIO-hM4Di-mCherry, hSyn-DIO-hM3Dq-mCherry, hSyn-DIO-rM3Ds-mCherry, hSyn-DIO-mCherry:* University of North Carolina Vector Core or AddGene) was pressure injected via Hamilton syringe (28ga) or glass micropipette bilaterally into the ventromedial midbrain over 2min (retracted 5min later) causing DREADD expression selectively in TH neurons of VTA (Coordinates relative to bregma: AP-5.5, ML+0.8, DV-8.15). WT rats were injected with one of the same DIO vectors (hM3Dq n=6, rM3Ds n=6, hM4Di n=5; Overall WT n=18), though no expression was observed in any rat with vectors from either source. Four+ weeks was allowed between virus injection and first CNO administration, during which time cocaine self-administration and extinction training occurred.

### Surgical Procedures

Intravenous (i.v.) catheter and intracranial surgeries are described in detail elsewhere(Mahler et al., 2012; Mahler et al., 2014). Chronic jugular i.v. catheters were implanted exiting the back, then intracranial virus injections were conducted stereotaxically in the same surgery. Cranial cannulae were implanted in a separate surgery after initial (8d) cocaine self-administration training (~3 weeks after the first surgery). After recovery, they completed 2 remaining self-administration days, followed by extinction, reinstatement tests, and food intake tests.

### Drugs

Cocaine HCl was dissolved in 0.9% saline (i.v. self-administration: 3.33mg/ml; i.v. behavioral economics self-administration: 2.73mg/ml; cocaine prime: 10mg/ml). CNO was obtained from Tocris or NIMH/NIDA Drug Supply Programs), dissolved in 5% DMSO in saline (0,1,10mg/ml), and injected i.p. 30min prior to behavioral tests. Yohimbine HCl (Sigma) was dissolved in water (2.5mg/ml) and injected i.p. immediately after vehicle/CNO injection. For intracranial injection, CNO was dissolved in 0.5% DMSO in ACSF (1mM) and microinjected 5min prior to behavior.

### Cocaine Self-Administration and Extinction

Operant testing was conducted in Med Associates chambers. Rats received initial cocaine self-administration training until they self-administered >10 infusions/daily 2hr session for 10d. AL presses yielded a 3.6sec infusion of 0.2mg/50µl i.v. cocaine and a concurrent tone/light cue, followed by a 20sec timeout period. Presses on the IL had no consequences. Following self-administration training, rats received at least 7d extinction training, when lever presses yielded neither cues nor cocaine, which continued until criterion was met (<25 AL presses for 2d).

### Cocaine Demand Elasticity

Rats (hM4Di: n=10; hM3Dq: n=6; rM3Ds: n=7; WT: n=8) were trained to self-administer cocaine under a within-session cocaine demand protocol as described previously(España et al., 2010; Oleson et al., 2011; Bentzley et al., 2013). On an FR1 schedule, rats self-administered cocaine at doses decreasing each 10min interval of the session (383.5, 215.6, 121.3, 68.2, 38.3, 21.6, 12.1, 6.8, 3.8, 2.2, and 1.2 µg per infusion), requiring increasing effort across the session to maintain preferred cocaine blood levels. After every session, an exponential demand equation(Hursh, 1993; Bentzley et al., 2013) was fit to the results from each rat to determine economic demand for cocaine as prevously described(Bentzley et al., 2013). Rats were trained for a minumum of 5d, until <25% varibility of the α parameter in the last 3 sessions. Therefore, effects of neural manipulations on 1) preferred cocaine blood levels under low effort conditions (free consumption; Q_0_), and 2) sensitivity of demand to price (demand elasticity; α) can be derived from data in a single session. Values for α and Q_0_ were calculated as percentage of stable performance on the prior 3 drug-free training days.

### Reinstatement

Following extinction training, rats underwent a series of 2h reinstatement tests, each separated by 2+ days of re-extinction. Each reinstatement modality was tested after counterbalanced vehicle/CNO(1,10mg/kg) injections, before moving on to the next reinstatement modality. The order of reinstatement modalities (**Fig.1**) was fixed to minimize potential carryover effects of one type of reinstatement (e.g. YOH or cocaine prime) on others (e.g. cue reinstatement)(Mahler et al., 2012). For cued reinstatement tests, AL presses yielded cocaine-associated 3.6sec-duration cue presentations (followed by a 20sec time out period signaled by the houselight turning off), but no cocaine. IL presses were recorded, but inconsequential throughout training and testing. For the extinguished context test, lever presses yielded neither cocaine nor cocaine cues, but were recorded. For primed reinstatement tests, i.p. cocaine (10mg/kg) was given immediately before the test and lever presses were also inconsequential. For yohimbine+cue reinstatement tests, rats received the pharmacological stressor yohimbine (2.5mg/kg i.p.; 5min before CNO/Veh, 35min before test) and AL presses yielded delivery of previously cocaine-associated cues, followed by a 20sec signaled timeout. After each reinstatement test, rats were given 2+ extinction re-training sessions, until they returned to the extinction criterion.

**Figure 1:**
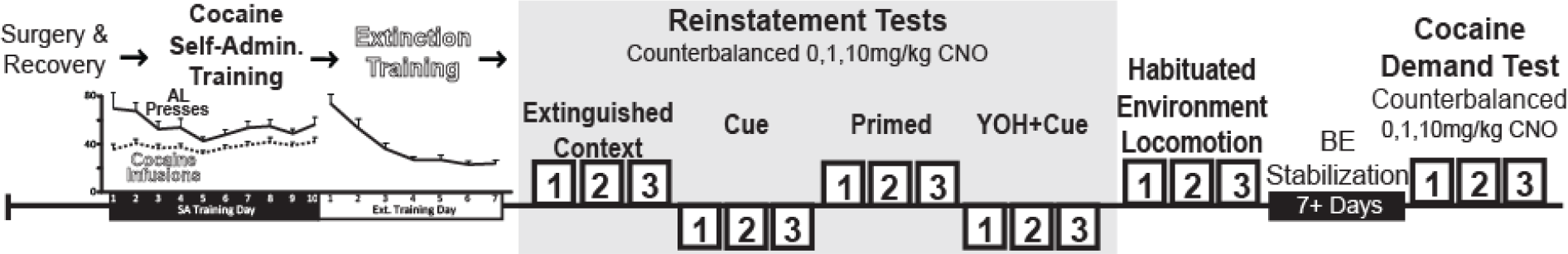
Behavioral Testing Timeline. Following surgery and recovery, Gq-, Gs-, Gi-DREADD rats and WT controls were trained to self-administer cocaine+ tone/light cue over 10 daily 2hr sessions, then extinguished over 7+ additional sessions. In a repeated measures design, we examined effects of counterbalanced injections of CNO (0,l,10mg/kg) on cocaine seeking in an extinguished context (no cues/cocaine), during reinstatement elicited by cocaine cues, cocaine priming injection, or yohimbine (YOH) +cocaine cues, on locomotor activity in a habituated environment, and on a test of cocaine economic demand.

### Locomotion

Following self-administration and reinstatement testing, rats (hM3Dq n=6, rM3Ds n=6, hM4Di n=5, WT n=7) were habituated for 2 daily 2hr sessions to a locomotor testing chamber (40×40×30cm), with clear plastic sides, corncob bedding, and photobeam arrays to measure horizontal and vertical activity, then tested for CNO effects over 2h.

### Food Intake

Effects of intracranial vehicle or CNO (1mM) injections into mPFC (n=10, 0.3µl), NAcCore (n=11, 0.5µl), or BLA (n=7, 0.5µl) on spontaneous chow intake were measured on two, 2hr sessions held 48 hours apart. Rats were first habituated for 2, 2hr sessions to the same clear plastic tub cage with corncob bedding, pre-weighed chow on the floor, and a water bottle. Chow intake was quantified by comparing food weight before and after these sessions.

### Co-Localization of mCherry and TH in VTA

To confirm DREADD expression in VTA DA neurons, co-staining for mCherry and TH was performed on VTA sections. 40µm sections were blocked in 3% NDS, then incubated in rabbit anti DS Red to label mCherry (Clontech Cat#632496, 1:1000) and mouse anti TH (Immunostar cat#22941; 1:1000) for 16hrs at RT. Sections were washed, then incubated in AlexaFluor donkey anti rabbit 488 and donkey anti mouse 594 antibodies (both 1:500; Jackson) for 4hrs. Sections were coverslipped and imaged on a Leica epifluorescent microscope with StereoInvestigator software (MicroBrightfield). Images of the entire VTA and SN were obtained from 3 planes in the z-axis at 10X magnification, stitched together using the StereoInvestigator virtual slice module, and collapsed into a flat maximum projection image of ventral midbrain for quantification by an observer blind to experimental condition. The percentage of mCherry neurons with or without somatic TH expression, and TH(+) neurons with or without co-expression of mCherry, were quantified bilaterally within the entire VTA and SN on 1 slice/rat (5.3 to 5-6mm caudal of bregma; hM3Dq n=8; rM3Ds n=8; hM4Di n=8; mCherry n=6).

### Fos Expression in VTA DA Neurons

To verify DREADD stimulation of VTA DA neurons, we examined Fos expression in VTA TH(+) neurons in a subset of rats (hM3Dq veh/10CNO n=2/3; rM3Ds n=2/2; hM4Di n=2/2). Animals were given i.p. vehicle or CNO, left undisturbed in their home cages for 180min, then perfused. Coronal 40µm sections were blocked in 3% normal donkey serum, then incubated in rabbit anti c-Fos (Millipore ABE457, 1:1000) and mouse anti TH for 16hrs at RT in PBST. Sections were washed, then incubated in AlexaFluor donkey anti rabbit 488 and donkey anti mouse 594 (1:500 of each; Jackson cat#715585150) for 4hrs. Sections were imaged at 20x and Fos was quantified in TH(+) and TH(-) cells in 3 bilateral sections/rat at about −5.2, −5.5, & −5.8 caudal of bregma. All TH(+) and Fos(+) cells were counted, and expression of Fos in TH(+) neurons of VTA and SN was quantified.

### In Vitro Electrophysiology

Brain slices were prepared as described previously(Riegel and Williams, 2008; Williams et al., 2014) from behaviorally-tested TH:Cre rats with mCherry-tagged Gi-, Gq-, or Gs-coupled DREADDs as described above. Briefly, brains were removed following rapid decapitation and placed in a vibratome (Leica) containing ice-cold ACSF solution (126 mM NaCl, 2.5 mM KCl, 1.2 mM MgCl2, 1.4 mM NaH2PO4, 25 mM NaHCO3, 11 mM D-glucose, 0.4 mM ascorbate, and 0.01 mM MK801). Horizontal slices (220 m) containing the VTA were prepared and stored in oxygenated ACSF containing 0.01 mM MK801 (95% O2–5%CO2; 34°C) until recording. During recording, sections were perfused at a flow-rate of 2 ml/min with oxygenated ACSF, at 33°C. DA neurons residing 150 m from the lateral or medial side of the terminal nucleus of the accessory optic tract were visualized with infrared-differential interference contrast optics to confirm mCherry fluorescence indicative of DREADD expression. Recordings were made using Multiclamp 700B amplifiers (Molecular Devices) and collected with AxoGraph X (AxoGraph Scientific), filtered at 1–2 kHz, and digitized at 2–5 kHz. Spontaneous AP firing was monitored using either whole-cell current-clamp or cell-attached recording configurations with 1–2 M pipettes filled with 115 mM K-methylsulfate, 20 mM NaCl, 1.5 mM MgCl2, 2.5 mM HEPES, 2 mM ATP, 0.3 mM GTP, and 0.1 mM EGTA (pH 7.3; 265–270 mOsm). Firing rates were evaluated in the cell-attached configuration. Only neurons displaying stable pacemaker firing (1-5Hz) for the duration of the experiment were included for analysis of firing rates in response to CNO.

### In Vitro Cyclic Voltammetry

Separate TH:Cre rats (hM3Dq n=8, rM3Ds n=8, hM4Di n=6, WT n=8) were killed, and their brains rapidly removed and prepared as previously described(Brodnik and España, 2015). Coronal slices (400µm) of the striatum were maintained at 32 °C in oxygen-perfused (95% O_2_–5% CO_2_) artificial cerebrospinal fluid, which consisted of (in mM): NaCl, 126; NaHCO_3_, 25; D-glucose, 11; KCl, 2.5; CaCl_2_, 2.4; MgCl_2_, 1.2; NaH_2_PO_4_, 1.2; L-ascorbic acid, 0.4; pH adjusted to 7.4. A carbon fiber microelectrode (150–200µm length × 7µm diameter) and a bipolar stimulating electrode (Plastics One) were placed in the NAc. The carbon fiber electrode potential was linearly scanned as a triangular waveform from −0.4 to 1.2 V and back to −0.4 V (Ag vs. AgCl) using a scan rate of 400 V/s. Cyclic voltammograms were recorded at every 100ms by means of a potentiostat (Dagan, Minneapolis, MN, USA) using Demon Voltammetry and Analysis(Yorgason et al., 2011). Extracellular concentrations of dopamine were assessed by comparing the current at the peak oxidation potential for dopamine with electrode calibrations with 3µM dopamine. Dopamine release was evoked every 5min by a single 4ms, stimulation (monophasic, 400µA), until three stable baseline responses were acquired (less than 10% variation) before CNO was added to the perfusion medium. Following application of CNO, we evaluated DA release to both single pulse and multiple pulses (five pulses at 5-40 Hz).

### Anesthetized Electrophysiology

In separate rats (n=22; hM3Dq n=8; rM3Ds n=2; hM4Di n=8; mCherry n=4), we performed extracellular, anesthetized recordings of single units in VTA, before and after i.p. injection of CNO (10mg/kg). Animals were anesthetized with 2-3% isoflurane and maintained at a rectal temperature of 36-38°C throughout recordings. Affixed in a stereotaxic instrument, craniotomies were prepared dorsal to VTA, exposing the brain surface from 0.5-1.3mm lateral of midline and 4.0-7.0 caudal of bregma. Following previously described procedures(Georges and Aston-Jones, 2002; Kaufling and Aston-Jones, 2015), glass micropipettes (filled with 2% pontamine sky blue in 0.5M sodium acetate; tip internal diameter ~1µm; impedance 6-12mΩ) were advanced through VTA between the following coordinates (relative to bregma): AP: −5.1-6.1; ML: +0.5-1.1, DV: −7.0-9.0, implemented in a consistent grid-based sampling method(Grace et al., 2007) with 200µm+ between consecutive tracks and <400µm between at least one pre-and post-CNO track within a given animal. Signals were amplified and filtered (1000x gain, 50Hz-4000Hz bandpass) using conventional electronics, sent to an audio monitor and oscilloscope for online monitoring (low pass: 50Hz, high pass: 16KHz), and digitized and recorded via CED 1401 and Spike 2 software (Cambridge Electronic Design). When single unit activity was detected by monitoring audio and oscilloscope outputs, we advanced slowly upon the cell until it was isolated at ~2:1 signal/noise ratio or better. All isolated units (n=400) were recorded for 120sec, then the electrode was lowered 200µm, and sampling resumed. After sampling 2-9 tracks/rat, we injected CNO (10mg/kg, i.p.), and from 30-210min later, we sampled an additional 1-4 tracks/rat (maximum 3h after CNO injection). Pontamine sky blue was deposited electrophoretically at the end of the last track each day prior to perfusion, and recording sites within VTA were reconstructed. Cells were classified as Type 1 (“Classic DA-like”) or Type 2, based on conventional criteria described in detail elsewhere(Grace and Bunney, 1984; Luo et al., 2008; Ungless and Grace, 2012). The mean number of Type 1&2 cells encountered per track was compared before and after CNO.

### Co-localization of mCherry and TH in mPFC and NAc

We examined co-localization of TH (a putative marker of DAergic axonal microdomains(Zhang et al., 2015)) with mCherry in cocaine-experienced rats described above. We imaged areas of prelimbic cortex (layers 5&6) and NAcCore (immediately dorsal and medial of the anterior commissure) from DREADD-infected rats (hM3Dq n=4; rM3Ds n=2; hM4Di n=4). Z-series confocal microscopy data sets were acquired with the exact same parameters for PFC and NAc sections. Images were taken through the entirety of the tissue slice for both regions using a 63X objective with a voxel size of 0.24 × 0.24 × 0.504. Two z-series data sets were taken for each animal, one on each hemisphere. Values obtained for these two data sets were averaged to create an animal average for each brain region investigated. All laser and gain settings were kept constant between animals and between brain regions. Once data sets were collected they were deconvolved using autoquant deblur and imported into Bitplane Imaris. Once in Imaris, threshold values for colocalization analysis of DREADD and TH signals were automatically set by the Imaris software colocalization module. Values set by Imaris for colocalization signal intensity thresholds were not significantly different for DREADD (red) or TH (green) signals in either the mPFC or NAc, nor did mCherry expression differ between DREADD groups.

### Experimental Design and Statistical Analysis

For behavioral and colocalization, and Fos analyses, mixed model and repeated measures ANOVAs with Bonforroni corrected t-test posthocs, or paired samples t-tests were employed as appropriate. When data were not normally distributed, nonparametric statistics (Mann Whitney U; Friedman’s) or Grennhouse-Geisser degree of freedom corrections were used. Cocaine economic demand was stabilized for 2+ days prior to each test, then change from this stable baseline after vehicle or CNO was computed and analyzed with repeated measures ANOVA. Pearson correlations were computed to examine the relationship between locomotor and reinstatement effects of CNO (Change from vehicle day horizontal locomotion and cue-induced cocaine seeking after 10mg/kg CNO). For in vitro physiology, spontaneous firing rate in the 30sec before or after CNO bath application (5µM, 4-6mins) was measured in 1-2 mCherry fluorescing VTA neurons per rat. For in vitro voltammetry, evoked DA was computed as percent of pre-CNO baseline, and mixed model ANOVA determined effects DREADD group X stimulation frequency. For in vivo physiology, the mean number of Type 1 and 2 neurons per recording track per rat, before versus after CNO, were compared using unpaired t-tests in each DREADD group. All experiments were conducted in at least 2 cohorts, boosting confidence in the reliability of reported effects.

## Results

In male tyrosine hydroxylase (TH):Cre rats or their wild-type (WT) littermates (250-400g) bred at MUSC, UCI, or Drexel, we targeted injections of Cre-dependent double inverted open reading frame (DIO) vectors to VTA, causing expression of mCherry-tagged hM3Dq (Gq-coupled), rM3Ds (Gs-coupled), or hM4Di (Gi-coupled) DREADDs, or mCherry in VTA DA neurons in TH:Cre rats, or no DREADD expression in WT rats (**Fig.2a**). Expression of the DREADD-fused mCherry tag was highly specific to VTA/medial SN TH(+) neurons in TH:Cre rats, as previously reported(Witten et al., 2011; Mahler et al., 2014), with over 97% co-expression of TH in mCherry+ cell bodies in ventral midbrain(Tye et al., 2013; Mahler et al., 2014). Overall, >70% of VTA TH(+) neurons co-expressed DREADDs, while only ~10% of lateral SN dopamine neurons expressed DREADDs (**Fig. 2b**). DREADD expression spanned nearly the entire rostro-caudal axis of VTA in these animals, typically with sparser expression in medial than in lateral aspects of VTA, and extended into the most medial aspects of substantia nigra pars compacta (**Fig.2b**). No significant differences in VTA infection rate of cell bodies was seen between Gi, Gs, and Gq-coupled vectors (no effect of DREADD group on co-localization percentage: F_2,23_=0.576, p=0.57), and no DREADD/mCherry expression was seen in WT control rats.

**Figure 2:**
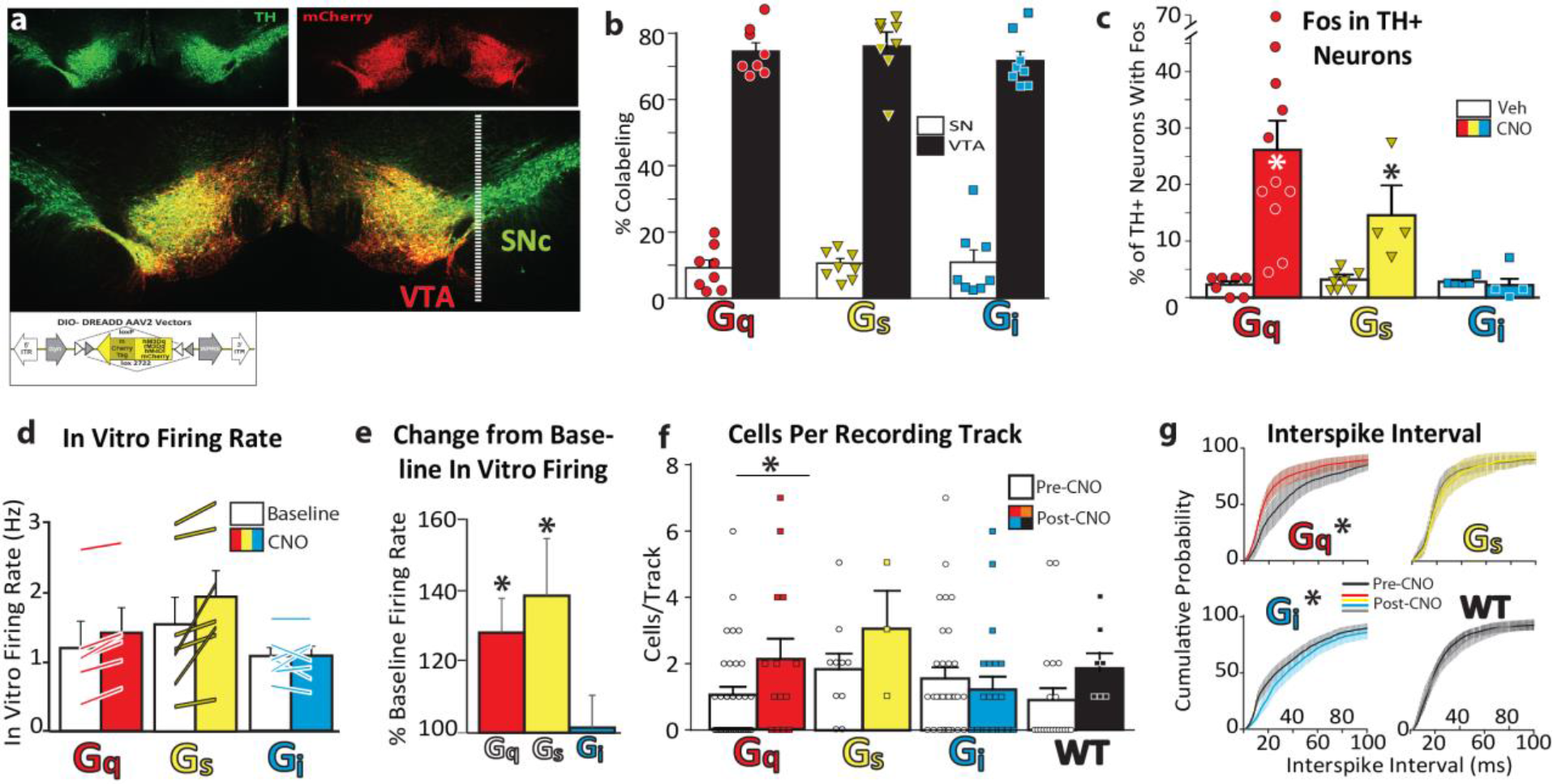
DREADD Expression and Function in VTA DA Neurons. **a.** Midbrain tyrosine hydroxylase (TH) staining (green) and mCherry-tagged DREADD expression (red) in VTA is shown, with yellow indicating co-labeling. Inset shows schematic of AAV double-inverted open reading frame (DIO)-DREADD vectors used. **b.** Percentage of midbrain TH+ cells located within borders of substantia nigra pars compacta (SN; white bars), or VTA (black bars). Individual rat data shown with red diamond (Gq), yellow triangle (Gs) or blue square (Gi) symbols, **c.** Percentage of TH+ VTA neurons expressing Fos after Vehicle or CNO (lOmg/kg) in Gq-, Gs-, or Gi-coupled DREADD rats (individual rat data represented with symbols on top of bars), **d.** Firing rates are depicted for baseline and post-CNO periods, individual cells depicted with lines, **e.** CNO increases mean (SEM) firing rate of VTA DREADD+ neurons in vitro, relative to baseline firing, **f.** In anesthetized rats with hM3Dq DREADDs, more Type 1 cells were encountered per electrophysiological recording track after 10mg/kg CNO (colored bars, symbols) than pre-CNO (white bars, symbols). No significant effects on population activity were observed after CNO in Gs, Gi, or WT groups, **f.** Interspike interval before CNO (black lines, error bars=SEM) and after 10mg/kg CNO *i.p.* (red=Gq, yellow=Gs, blue=Gi, black=WT). *p<0.05.

As expected, CNO (10mg/kg) markedly induced Fos in TH(+) VTA neurons of rats expressing hM3Dq, but not hM4Di DREADDs (DREADD group X treatment (Tx) interaction; F_2,35_=3.4, p=0.046; **Fig.2c**). Acute stimulation of VTA DA neurons with CNO (10mg/kg) in hM3Dq rats increased over vehicle the percentage of TH(+) VTA neurons that expressed Fos (main effect of Tx in hM3Dq rats; Mann-Whitney U (MWU) test: p=0.001), whereas Fos in TH(-) VTA neurons was concurrently suppressed (MWU p=0.03; **Fig.2c**). In rM3Ds rats, CNO similarly increased Fos in TH(+) neurons (Tx effect; MWU p=0.004), but Fos expression in TH(-) VTA neurons was not altered (MWU: p=0.68, n.s.). Fos was not significantly reduced by CNO in hM4Di rats [TH(+) neurons Tx effect: MWU, p=0.26, n.s.; TH(-) neurons: p=1.0, n.s.; **Fig.2c**], likely because Fos expression was already very low in control rats treated with vehicle in their home cage.

To further characterize DREADD modulation of DA cell activity, we recorded effects of CNO on firing in vitro and in vivo. First, horizontal VTA slices were prepared as described previously(Williams et al., 2014), and DREADD-expressing (mCherry+) cells were sampled in the cell-attached configuration to record spontaneous firing rates. We detected no difference in the basal firing rates of DA neurons expressing hM3Dq, rM3Ds, or hM4Di (Hz: F_2,16_=0.658, p=0.531). After washing on CNO (5µM), both hM3Dq and rM3Ds-expressing DA neurons increased firing compared to pre-CNO baseline (hM3Dq: t_6_=2.9, p=0.028; rM3Ds: t_4_=5.4, p=0.006), but CNO did not affect spontaneous firing in slices from hM4Di rats (t_6_=0.04, p=0.968; **Fig.2d-e**).

We next examined whether CNO affects the number of spontaneously active VTA cells recorded *in vivo* in isoflurane-anesthetized rats. Using a systematic grid-based cells/track sampling method(Lodge and Grace, 2006), we recorded all well-isolated cells encountered in the 3.5 hours before and after i.p. CNO injections (n=257 pre-CNO; n=143 post-CNO). We analyzed data from the two most commonly sampled types of neurons encountered in VTA: Type 1 neurons with >2ms waveform and <10Hz firing rate, which are likely to be a subset of DAergic neurons(Ungless and Grace, 2012) (n=200), and Type 2 neurons with <2ms waveforms, >10Hz firing rate, and an uncharacterized phenotype (n=200). CNO (10mg/kg) did not affect the numbers of Type 1 or Type 2 cells encountered in hM4Di, rM3Ds, or mCherry rats (hM4Di: F_1,49_=0.3, p=0.57; rM3Ds: F_1,12_=1.3, p=0.27; mCherry: F_1,26_=3.1, p=0.09; Type 2: Fs<0.73, ps>0.4), but did increase prevalence of active Type 1 cells in hM3Dq rats (F_1,46_=4.0, p=0.05) without affecting Type 2 prevalence (F_1,46_=0.72, p=0.790). This indicates that Gq stimulation induced activity in a subset of otherwise quiescent VTA DA cells (**Fig.2f**). We also examined effects of DREADD manipulations on firing rates of VTA DA neurons and found, relative to interspike intervals (ISIs) prior to CNO, CNO decreased ISIs in hM3Dq rats, increased it in hM4Di rats, and did not alter ISI in rM3Ds or WT rats (hM3Dq: F_(1,2000)_=125.5, p<0.0001; hM4Di: F_(1,3600)_=57.45, p<0.001; rM3Ds_(1,800)_=1.85, p=0.1737; mCherry: F_(1,1600)_=2.03, p=0.1543; **Fig.2g**). Having validated chemogenetic modulation of DA neurons, we next examined effects of these manipulations on cocaine seeking behaviors.

### Dual Roles for DA Neurons in Economic Demand for Cocaine

We first queried whether chemogenetic DA neuron modulation affects the free consumption of, or motivation for, cocaine. Across species, motivation for a reward can be measured via analysis of demand elasticity, or the sensitivity of reward consumption to increasing price(Bickel et al., 1990; Hursh, 1993). A useful behavioral tool for examining effects of acute neural manipulations in rats on economic demand is the within-session cocaine behavioral economic (BE) paradigm(España et al., 2010; Oleson et al., 2011; Bentzley et al., 2013; Bentzley et al., 2014). Rats were trained on 10 daily 120min sessions to press an active lever (AL) (not an inactive lever; IL) to receive cocaine (0.2mg/infusion, i.v.) accompanied by a tone/light cue, as described below. They were then trained on 110 min BE sessions, wherein cocaine was delivered under an increasing effort requirement, allowing within-session analysis of cocaine demand elasticity. Data were fit to an exponential demand curve(Hursh and Silberberg, 2008) and the demand parameters Q_0_ (free consumption) and α (demand elasticity; inverse of motivation) were derived as previously described(Bentzley et al., 2013). Because α and Q_0_ are stable across consecutive days of training, these values can also be compared across sessions, revealing effects of a neural manipulation on cocaine demand (Oleson et al., 2011; Bentzley et al., 2014).

CNO in hM3Dq rats robustly decreased free cocaine consumption (Tx effect on Q_0_, F_2,12_=6.584,p=0.012; **Fig.3b**), indicating that increased Gq signaling in these neurons partially substitutes for cocaine’s subjective effects. Interestingly, demand elasticity (α parameter) in hM3Dq rats also decreased, indicating that DA neuronal activation increased effort expended to obtain cocaine as doses decreased during the session, indicating increased motivation for drug (Tx effect on α; F_2,12_=11.44, p=0.002; **Fig.3a**). In contrast, stimulation of rM3Ds DREADDs in DA neurons failed to significantly affect either free consumption or demand elasticity (α: F_2,14_= 0.03217, p=0.97; Q_0_: F_2,14_= 0.83, p=0.46). Stimulation of hM4Di DREADDs in VTA DA neurons caused animals to consume more cocaine under low effort conditions (Tx effect on Q_0_: F_2,24_=3.771, p=0.038), and demand became more elastic (α increased, motivation decreased; Tx effect on α: F_2_,_2_4= 4.42, p=0.023); opposite to Gq-stimulation effects. No effects of CNO were observed in WT control rats (α: F_2,14_=1.765, p=0.207; Q_0_:F_2,14_=0.009, p=0.991; individual rat data: **Extended Fig.3-1**). These results together indicate that DA neurons play at least two distinct roles in cocaine intake—mediating the subjective effects of cocaine under low effort conditions and facilitating motivation to pursue cocaine under high effort conditions. Future studies should determine whether the same DA neurons underlie functions related to cocaine free consumption and motivation, or whether these are instead mediated by separate DA neurons within VTA.

**Figure 3:**
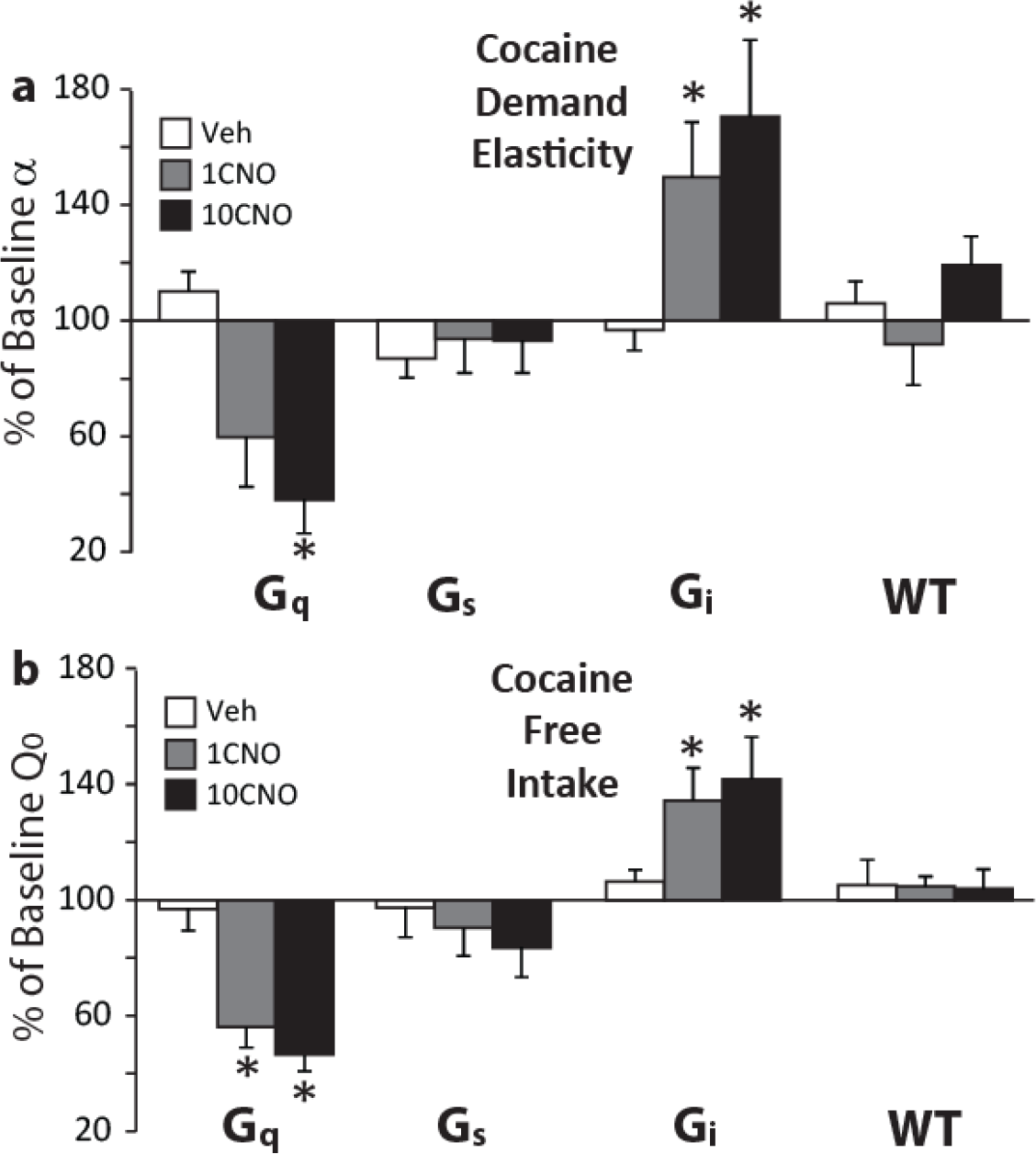
DA Neuron Modulation Affects Cocaine Free Intake and Demand Elasticity. **a.** Effects of CNO (0,1 or 10mg/kg) in Gq, Gs, Gi DREADD, or WT rats on demand elasticity (a) are depicted, b. Effects of CNO on cocaine free intake (Qo). When VTA DA neurons were stimulated, rats preferred less cocaine but worked harder to obtain this lower dose. When DA neurons were inhibited, subjects preferred more cocaine but were less willing to work hard to obtain it. *p<0.05. Individual rat data shown in **Extended Fig. 3-1**

### DA Neuron Roles in Cocaine Reinstatement

Addiction is a chronic relapsing disorder, so we examined chemogenetic DA neuron manipulation effects in a rat model of relapse elicited by cocaine cues, priming, or stress(Bossert et al., 2013; Martin-Fardon and Weiss, 2013). Although these stimuli elicit similar levels of drug seeking behavior, the neural substrates underlying each type of reinstatement are distinguishable(Stewart, 2000; Kalivas and McFarland, 2003; Mahler et al., 2014). Therefore, we sought to determine the necessity and sufficiency of VTA DA neurons for reinstatement of cocaine seeking.

After 10d cocaine self-administration followed by 7d+ extinction training, we queried effects of DA neuron stimulation or inhibition on reinstatement behavior (i.e., pressing the previously cocaine-delivering AL) due to 1) DA neuron manipulation alone (CNO in DREADD-expressing rats in an extinguished context without cues or cocaine), 2) cue-induced reinstatement (AL response-contingent cocaine cues, but no cocaine), 3) cocaine-primed reinstatement (10mg/kg cocaine i.p. prior to test, no cues or additional cocaine), and 4) pharmacological stress enhancement of cued reinstatement [2.5mg/kg i.p. yohimbine HCl (YOH), with response contingent cues but no cocaine]. Each reinstatement modality was tested three times in each rat, each 30min after counterbalanced injections of CNO (0, 1 & 10mg/kg; **Fig.1**).

### Gq DA Neuron Stimulation Dramatically Increased Reinstatement

Under all reinstatement conditions examined, CNO robustly increased cocaine seeking behavior up to 2000% of vehicle day levels in hM3Dq rats (effect of Tx on: extinguished context: F_2,12_=4.8, p=0.03; cued reinstatement: F_2,12_=6.1, p=0.02; primed reinstatement: F_2,12_=7.1, p=0.009; YOH reinstatement: F_2,12_=4.1, p=0.045; **Fig.4a**, **Extended Fig. 4-1**). CNO effects were specific to the AL (as opposed to the always inert IL) in the extinguished context (Tx X lever interaction: F_2,12_=6.2, p=0.014), and AL specificity also approached significance in cue reinstatement (F_2,12_=3.4, p=0.068). CNO increased pressing on both levers equivalently on cocaine-primed (F_2,12_=1.7, p=0.23) and YOH reinstatement tests (no interaction: F_2,12_=2.2, p=0.158). CNO effects on AL pressing did not statistically differ by CNO dose (change from vehicle day AL pressing after 1 or 10mg/kg CNO; ts<1.55, ps<0.171), nor did effects dissipate over the 2h test (no dose X time interaction: Fs_6_,36<2.4, n.s.; **Fig.5a**).

**Figure 4:**
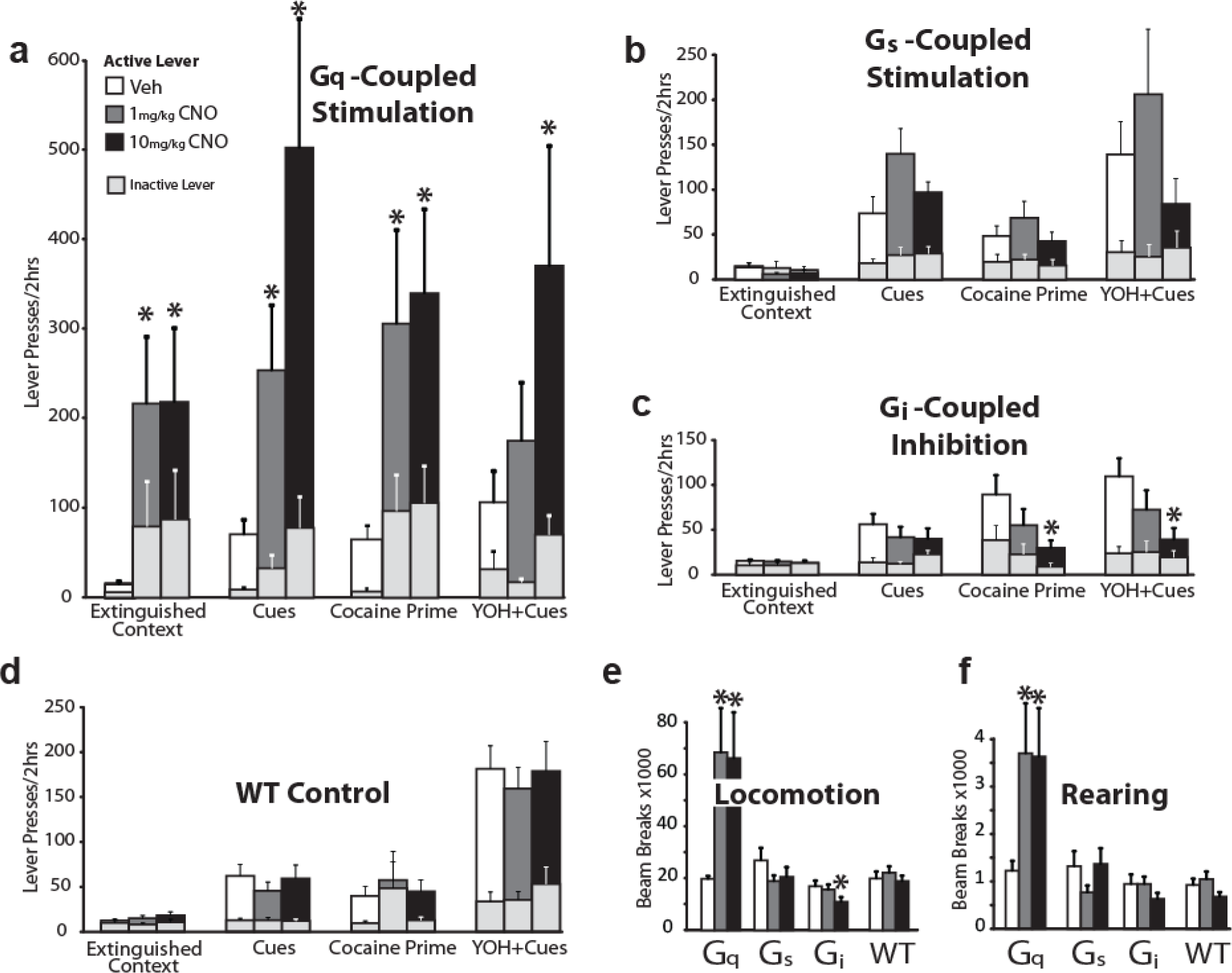
Reinstatement of Cocaine Seeking is Modulated by Chemogenetic DA Neuron Manipulations. **a.** In Gq-DREADD rats, CNO (1mg/kg: grey bars, 10mg/kg: black bars) increased cocaine seeking (relative to vehicle: white bars) in the absence (extinguished context) or presence of cues, and after a cocaine prime (no cues) or yohimbine (YOH+cues) test (10mg/kg). **b.** CNO failed to increase reinstatement in rM3Ds Gs-DREADD rats. **c.** In Gi-DREADD rats, CNO reduced the priming effects of cocaine and potentiation of cued responding by YOH (10mg/kg). **d.** No effects of CNO were observed in WT rats with no DREADD expression. Gq-stimulation increased, and Gi-stimulation decreased locomotor activity in a familiar environment. Light grey bars indicate IL presses for each test. *p<0.05 e. Effects of CNO (0,1,10mg/kg) on horizontal locomotion (beam breaks) and **f.** rearing behavior is shown for DREADD-expressing and WT animals. Individual rat data shown in **Extended Fig. 4-1**.

**Figure 5:**
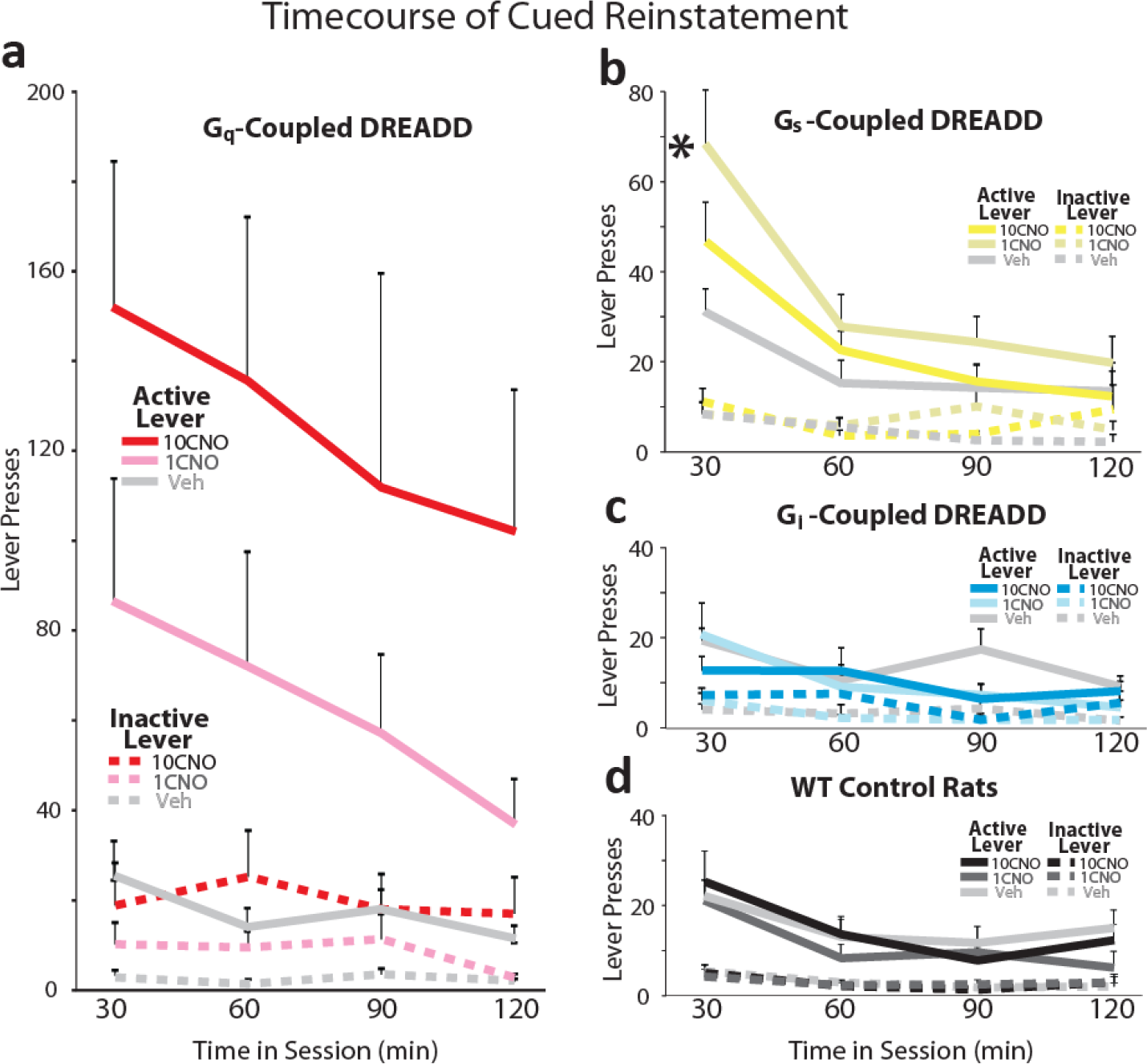
Timecourse of Cue-Induced Reinstatement. Timecourse of AL (solid lines) and IL (dashed lines) are shown for **a)** hM3Dq, **b)** rM3Ds, **c)** hM4Di, and **d)** WT control rats after vehicle or CNO (1, 10mg/kg). Data are shown in 30min bins throughout the 120min session. Lines represent mean, error bars represent SEM. *p<0.05, interaction of dose X time.

### Modest Effects of Gs DA Neuron Stimulation on Reinstatement

In contrast to hM3Dq rats, CNO did not robustly enhance cocaine seeking in rats with equivalent expression of rM3Ds in VTA DA neurons (**Fig.4b**). Cued reinstatement was marginally increased by CNO (main effect of Tx: F_2,20_=4.780, p=0.02), but posthoc analysis of CNO dose effects on AL pressing failed to meet Bonferroni-corrected threshold (1mg/kg: t_10_=2.469, p=0.033; significance threshold: p<0.025). However, cue-induced AL pressing was increased by 1mg/kg (not 10mg/kg) CNO for the first 30min of the 2hr session, after which cocaine seeking returned to vehicle-equivalent levels (Tx X time interaction: F_2.4,23.7_=9.5, p=0.001; no effect on IL pressing: F_6,60_=0.9, p=0.5; **Fig.5b**). Other reinstatement modalities were not reliably affected by CNO in rM3Ds rats (extinguished context: F_2,16_=0.6, p=0.56; prime: F_2,18_=1.0,p=0.38; YOH: F_2,8_=1.6, p=0.26).

### Modality-Specific Effects of Gi-mediated DA Neuron Inhibition on Reinstatement

Effects of stimulating hM4Di receptors in VTA DA neurons on reinstatement behavior was notably dependent on the type of reinstatement tested, presumably because DA neurons are differentially recruited during cocaine seeking under these different behavioral circumstances (**Fig.4c**). In hM4Di rats, CNO attenuated reinstatement elicited by either a cocaine priming injection (F_2,14_=6.6, p=0.009) or YOH (F_2,14_=3.87, p=0.046). Pressing on both levers was decreased during primed reinstatement (no interaction of Tx X lever: F_2,14_=0.96, p=0.4), but only on the AL during YOH reinstatement (F_2,14_=4.5, p=0.031). In an extinguished context without cues, cocaine, or YOH, CNO failed to affect the already low levels of lever pressing (F_2,14_=0.04, p=0.97).

Intriguingly, despite the clear role for DA in conditioned seeking of cocaine and other rewards(Berridge, 2007; Everitt et al., 2008; Clark et al., 2012; Salamone and Correa, 2012), CNO in hM4Di rats did not significantly reduce cue-induced AL pressing relative to vehicle day (no Tx effect on AL pressing F_2,i4_=0.5, p=0.61). Instead, CNO (10mg/kg) in these animals *increased* pressing on the IL (F_2,18_=3.7, p=0.045; 1CNO: p=0.7; 10CNO: p=0.048), despite the fact that presses on this lever were never reinforced at any point in training or testing. After vehicle, rats pressed the AL 6 times more in response to cues than the IL, but this AL/IL ratio was decreased to ~2:1 after 10CNO (ratio of AL/IL presses: Veh: m=6.08+1.25; 1CNO: m=3.54+0.88; 10CNO: m=2.17+0.65: F_2,14_=5.7, p=0.015; 10CNO: t_7_=2.98, p=0.02; **Fig.6a**). Looking at responses across the session, CNO-treated rats performed similarly to vehicle day for the first 30min of the session, with predominant pressing on the cue-delivering AL(Mahler and Aston-Jones, 2012; Gipson et al., 2013). Following this initial 30min period, CNO-treated hM4Di rats began IL pressing more than on vehicle day (Tx X time interaction for AL/IL ratio: F_3,21_=3.15, p=0.046; **Fig.6b**). No effect of CNO on the AL/IL ratio was observed during other types of reinstatement in hM4Di rats or for other DREADD groups during reinstatement of any type (FS_2,14_<0.78, ps>0.48).

**Figure 6:**
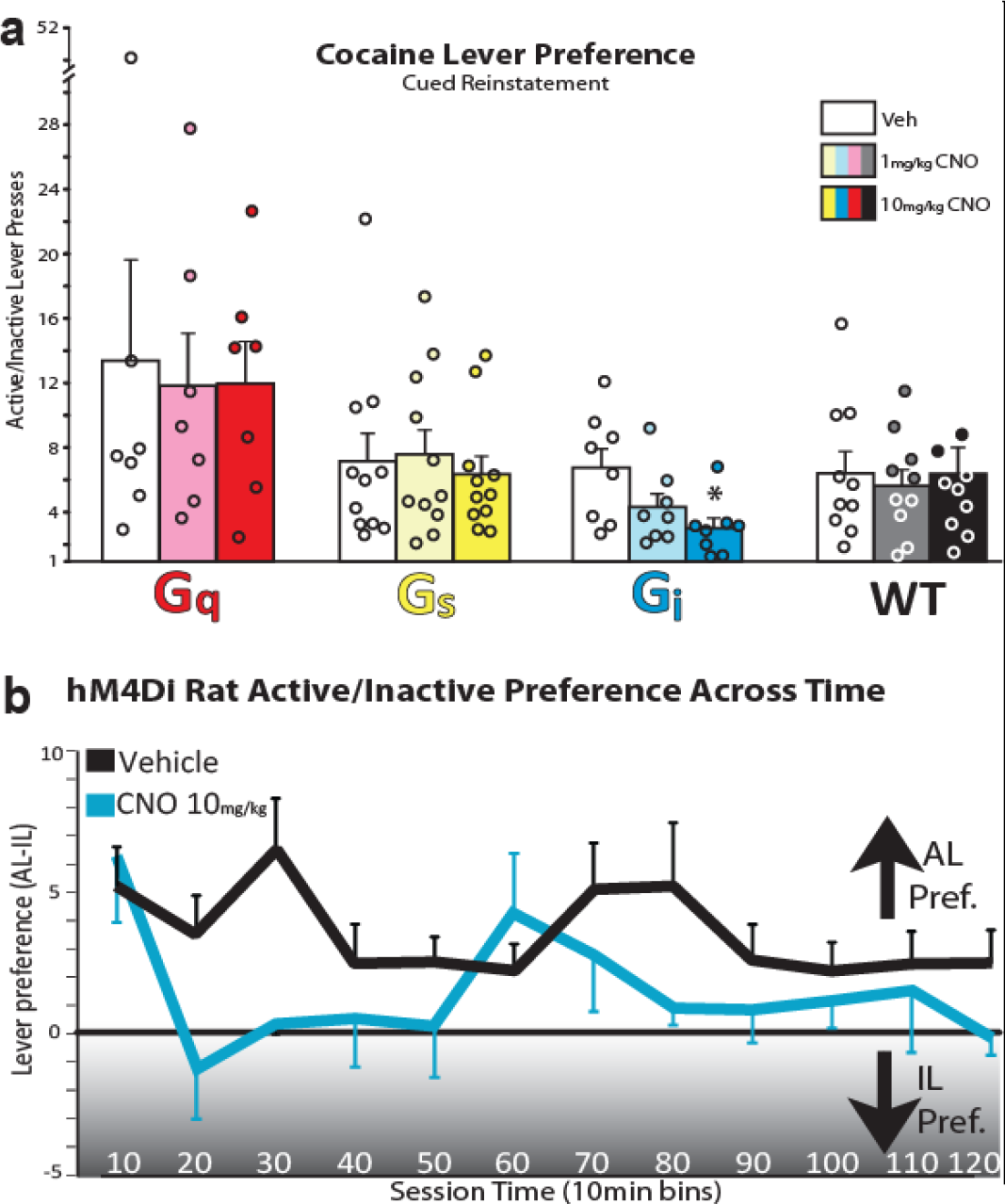
Stimulation of Gi-DREADDs in VTA DA Neurons Alters Cocaine Seeking Strategy. **a.** After vehicle (white bars) or CNO in Gq-DREADD, Gs-DREADD, or WT groups, rats pressed many times more often on the cue-delivering AL than on the always inert IL. In Gi-DREADD rats, CNO (lOmg/kg) instead decreased this active lever preference, **b.** Relative preference for the active over the IL is depicted in lOmin bins over the course of the 120min cue reinstatement test following vehicle (black line) or CNO (lOmg/kg, blue line). Initially (first lOmin), all rats predominantly pressed the AL, but as the session progressed, CNO receiving rats also underwent periods in which pressing was shifted to the IL instead. *p<0.05

### Validation of Presynaptic DA Release in NAc

DREADDs are robustly anterogradely transported, and readily observed in axons in forebrain DA neuron targets including NAc, medial prefrontal cortex (mPFC; **Fig.7a**), and basolateral amygdala (BLA). To verify chemogenetic modulation of presynaptic DA release, we prepared coronal slices of NAc from TH:Cre rats with hM3Dq, rM3Ds, or hM4Di-expressing VTA DA neurons, and DA release was measured using fast scan cyclic voltammetry(Ferris et al., 2013; Brodnik and España, 2015; Bucher and Wightman, 2015) before and after washing CNO (1µM) onto the slice, and in response to a single electrical pulse or multiple pulses across tonic and phasic frequencies (5 pulses at 5-40 Hz). CNO increased evoked DA in hM3Dq rats (F_1,21_=5.472, p=0.029), and decreased it in hM4Di rats (F_1,21_=12.77, p=0.0018), but did not affect release in rM3Ds or WT rats (F_1,21_=0.153, p=0.6989; **Fig.7c**). This indicates that DREADDs can be used to bidirectionally modulate DA release from VTA DA neuron terminals in a pathway-specific manner.

**Figure 7:**
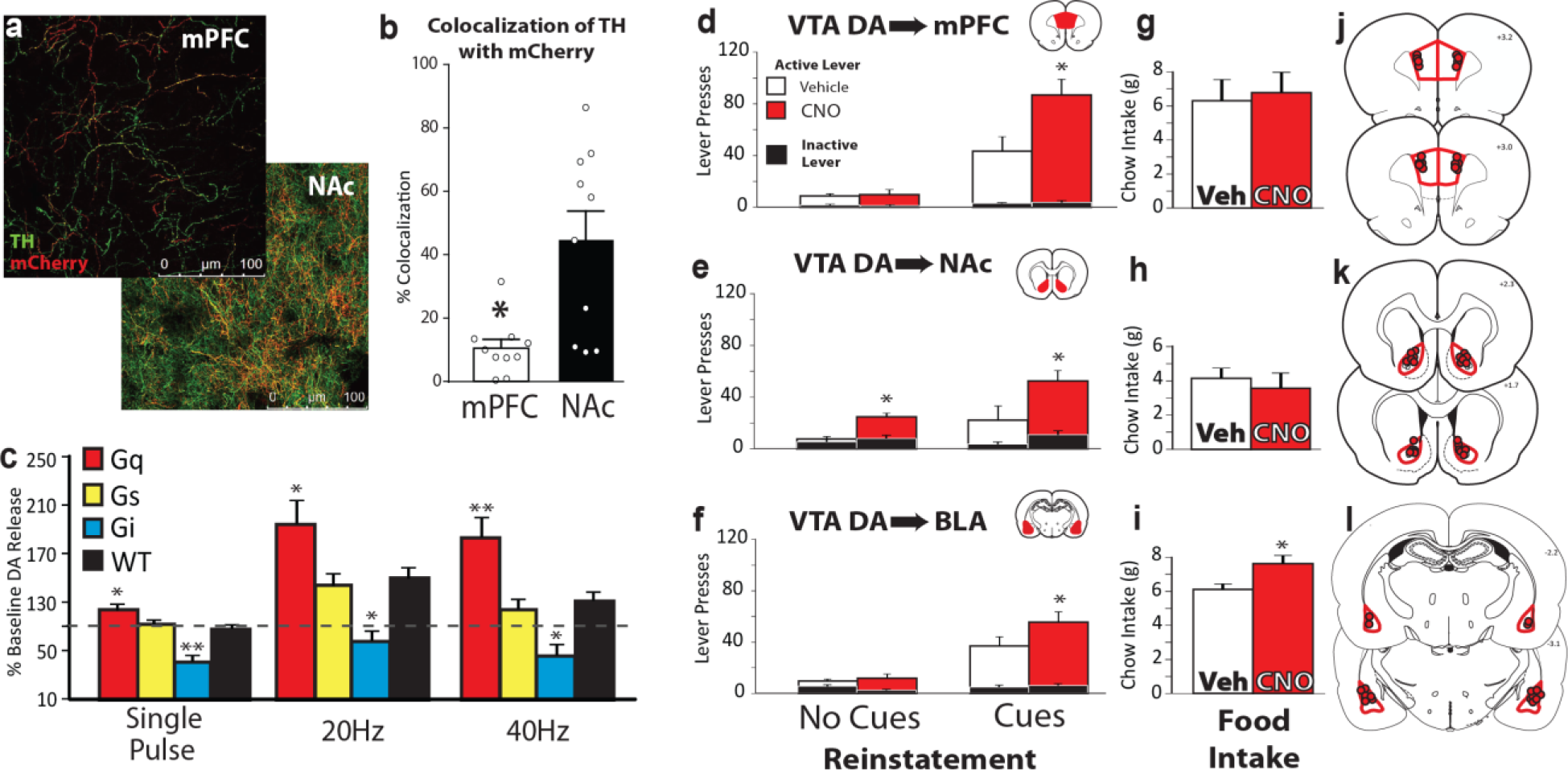
Chemogenetic Pathway-Specific Stimulation of VTA DA Terminals. **a.** hM3Dq DREADDs are robustly transported to targets including medial prefrontal cortex (mPFC) and nucleus accumbens (NAc). **b.** VTA TH(+) neuron terminals (labelled with mCherry (red) co-localized with tyrosine hydroxylase (TH; green) to a greater extent in NAc as compared to mPFC. **c.** In acute coronal NAc slices from rats with hM3Dq DREADDs in VTA DA neurons, DA release was evoked with local electrical stimulation (single pulse, 20Hz, or 40Hz) before or after bath application of CNO (1μM), measured using cyclic voltammetry. Bars depict *%* baseline (pre-CNO) DA release after CNO in Gq-, Gs- or Gi-DREADD rats, or WT rats. CNO facilitated DA release in Gq-DREADD rats at all stimulation parameters, and decreased DA release in Gi-DREADD rats. No effects of CNO were seen in Gs-DREADD or WT rats. Effects of CNO microinjection into mPFC **(d,g)**, NAc **(e,h)**, or BLA **(f,i)** on cocaine seeking in an extinguished context with (right bars) or without cues (left bars), or on 2hr chow intake in a separate test with the same rats **(j,k,l)**. Microinjection sites in j. mPFC, **k**. NAc, or **l**. BLA are depicted with red circles. *p<0.05, **p<0.01. Individual rat behavioral data shown in **Extended Fig. 7-1**.

### Efferent Target Specificity of Gq DREADD Stimulation Effects on Reinstatement

Given robust enhancement of reinstatement by hM3Dq DA neuron stimulation, we next sought to determine the corresponding contributions of VTA DA neuron projections to mPFC, NAc, and BLA for reinstatement or food intake behaviors. We and others have shown that applying CNO directly into the terminal fields of hM4Di-expressing neurons inhibits axonal release of neurotransmitters(Mahler et al., 2014; Stachniak et al., 2014; Wang et al., 2015; MacLaren et al., 2016), and here we examined for the first time whether projection-specific *stimulation* of dorsal mPFC, NAcCore, or BLA VTA DA neuron efferents affects behavior. A separate group of TH:Cre rats with DIO-hM3Dq in VTA and chronic bilateral cannulae in mPFC, NAc, or BLA (**Fig.7j,k,l**) were trained to self-administer cocaine for 10d, then extinguished of this behavior. Effects of CNO microinjections (vs. vehicle) on cocaine seeking in the absence or presence of response-contingent cues (extinguished context test or cued reinstatement test) were measured (**Fig. 7d-f**). CNO injections in any of the three structures increased AL pressing during cued reinstatement, indicating similarly increased impact of cues after stimulation of each DA neuron projection (mPFC: F_1,8_=14.024, p=0.006; NAc: F_1,12_=5.429, p=0.038; BLA: F_1,8_=6.424, p=0.035; **Fig. 7d-f**, **Extended Fig.7-1b**). Increased pressing was specific to the AL for mPFC and BLA, and approached significance for NAc (Tx X lever interaction: mPFC: F_1,8_=8.676, p=0.019; NAc: F_1,12_=3.887, p=0.072; BLA: F_1,8_=8.26, p=0.021). Importantly, NAc CNO, but not mPFC or BLA CNO, also *induced* cocaine seeking in the absence of cues, cocaine, or YOH (Tx effect on AL pressing in an extinguished context; mPFC: F_1,7_=0.104, p=0.757; NAc: F_1,12_=15.448, p=0.002; BLA: F_1,8_=0.019, p=0.894; **Fig. 7d-f**; **Extended Fig. 7-1a**). This shows that stimulation of VTA DA terminals in NAc alone is sufficient to generate seeking behavior even in the absence of endogenous circuit activity induced by Pavlovian cues. In the same rats, we next examined food intake for 2 hours after intra-mPFC, NAc, or BLA Veh/CNO microinjections (**Fig. 7d-f**, **Extended Fig. 7-1c**). Neither mPFC nor NAc CNO injections altered spontaneous eating (mPFC: t_9_=0.312, p=0.762; NAc: t_10_=0.782, p=0.452), but BLA CNO increased food intake (t_6_=2.466, p=0.049)—to our knowledge, the first demonstration of unconditioned feeding induced by a manipulation of BLA. These findings show novel roles for DA neuron efferents to NAc in cocaine seeking, and to BLA in unconditioned food intake.

### Colocalization of Axonal DREADDs with TH

Though DA neuron axonal stimulation had behavioral effects consistent with enhancement of DA release, and such enhanced DA release is observed in NAc (**Fig.7c**), DA neurons release other transmitters besides DA, including glutamate(Sulzer et al., 1998; Lapish et al., 2006; Hnasko et al., 2012; Trudeau et al., 2014; Zhang et al., 2015; Barker et al., 2016) and GABA(Tritsch et al., 2012; Stamatakis et al., 2013; Root et al., 2014). Perhaps relatedly, a marked lack of co-localization of mCherry [expressed exclusively in TH(+)neurons in VTA(Mahler et al., 2014)] and TH (a requisite enzyme for DA production(Zhang et al., 2015)) was observed in some targets, especially in mPFC. There was a significantly higher percentage of DREADD signal (red) colocalized with TH terminals (green) in NAc when compared to the mPFC (t_18_=3.530; p=0.0024; **Fig.7b**). Further inquiry into the neurochemical identity and behavioral functions of these TH-negative terminals originating from mPFC DA neurons is warranted, particularly given recent reports of marked neurochemical heterogeneity within VTA DA neurons and their axons(Tritsch et al., 2012; Trudeau et al., 2014; Zhang et al., 2015; Barker et al., 2016). *Specificity and Reversibility of DREADD Stimulation Effects:* Though CNO can have off-target effects affecting behavior(MacLaren et al., 2016; Gomez et al., 2017), effects here were specific to DREADD-expressing rats as there was no effect of either CNO dose on: lever pressing in WT rats, lever pressing during the extinguished context test (F_2,18_=1.13, p=0.344), cue (F_2,18_=0.73, p=0.495), primed (F_2,18_=1.42, p=0.27) or YOH reinstatement (F_2,16_=0.65, p=0.537; **Fig.4**), or cocaine demand (α: F_2,14_=1.765, p=0.207; Q0:F_2,14_=0.009, p=0.991; **Fig.3**). Neither did we find evidence of constitutive effects of DREADDs in the absence of CNO, as behavior of hM3Dq, rM3Ds, hM4Di, and WT rats did not differ during cocaine self-administration (total self-administered cocaine infusions; Fs_3,32_<1.8, ps>0.17; **Fig.8a-b**) or lever pressing during extinction (no day X group interaction: F_8.7,92.8_=1.6, p=0.14; **Fig.8c**), nor did groups differ on vehicle day in reinstatement behavior (extinguished context: F_3,30_=0.189, cue: F_3,32_=0.27, prime: F_3,31_=2.238, YOH: F_3,25_=1.697; **Fig.8d**), cocaine demand (a: F_3,28_=1.055, p=0.3839), or locomotor activity (Horiz F_3,29_=1.728; Rears: F_3,29_=0.664).

**Figure 8:**
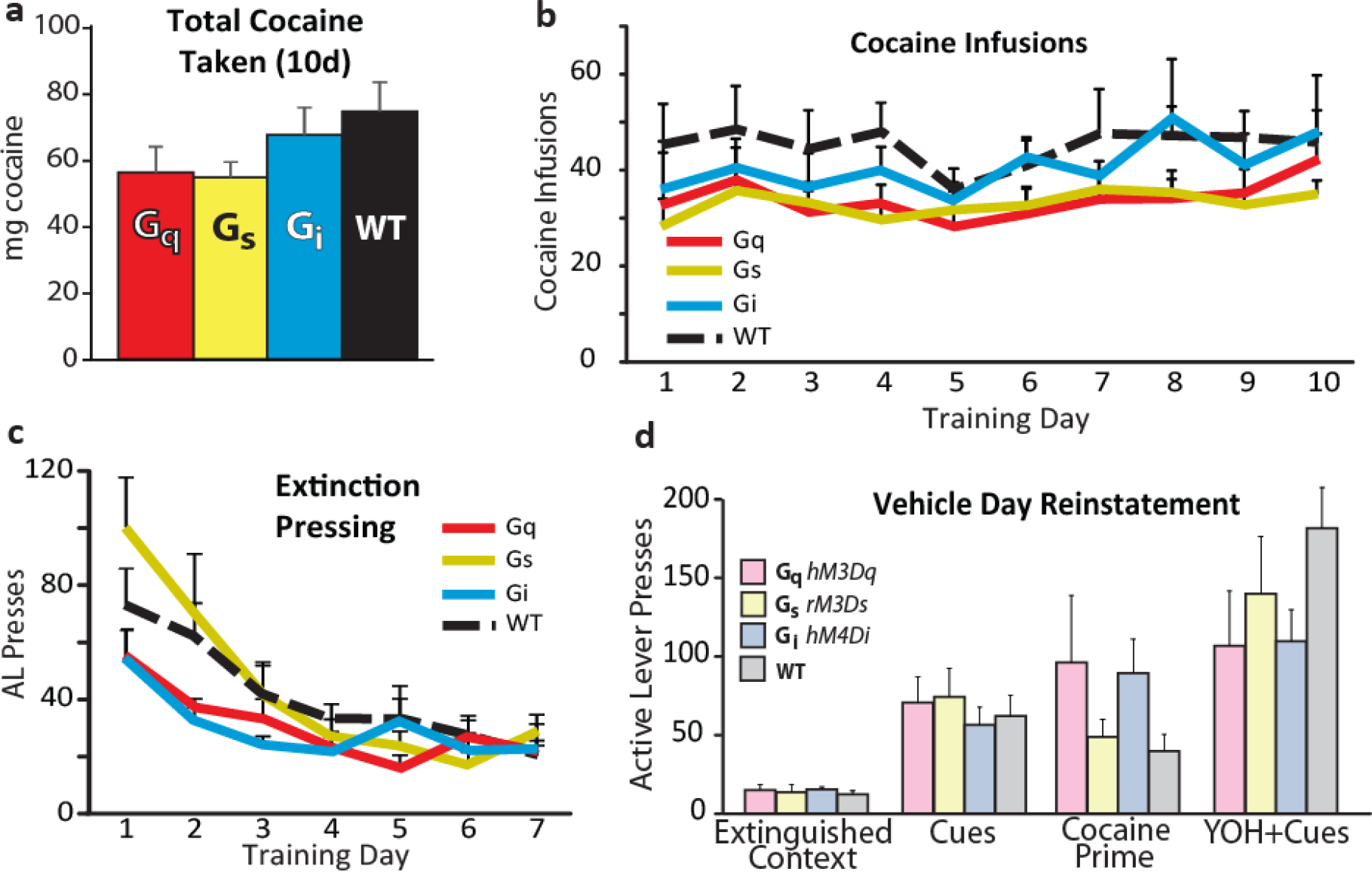
DREADDs do not Affect Behavior in the Absence of CNO. No significant effects of DREADD expression are observed in the absence of CNO. Rats with DREADDs (Gq=red, Gs=yellow, Gi=blue) or no DREADDs (WT controls=black) did not differ in **a)** the total mg cocaine self-administered throughout training, **b)** the number of daily cocaine infusions across the 10d training, **c)** AL pressing across the first 7d of extinction training, or **d)** AL pressing on reinstatement. Bars and lines represent means, error bars represent SEM.

Given the intimate involvement of G-protein signaling in synaptic plasticity, and prior reports of persistent effects of rM3Ds DREADD stimulation(Ferguson et al., 2013; Nakajima et al., 2016), we searched for persistent effects of prior-day CNO (compared to prior-day Veh) on behavior. No such effects were found during extinction retraining days following each CNO/vehicle reinstatement test (extinction test: F_2,16_=0.141; cue: F_2,20_=0.331; prime: F_2,18_=0.57; YOH: F_2,50_=0.078), or on cocaine intake following CNO/vehicle economic demand tests (no effect of Tx on α: F_2,36_=1.229, or Q_0_: F_2,36_=0.846; **Fig.9**). Together, these findings show that DREADD manipulations specifically and reversibly increase or decrease VTA DA neuron function as expected.

**Figure 9:**
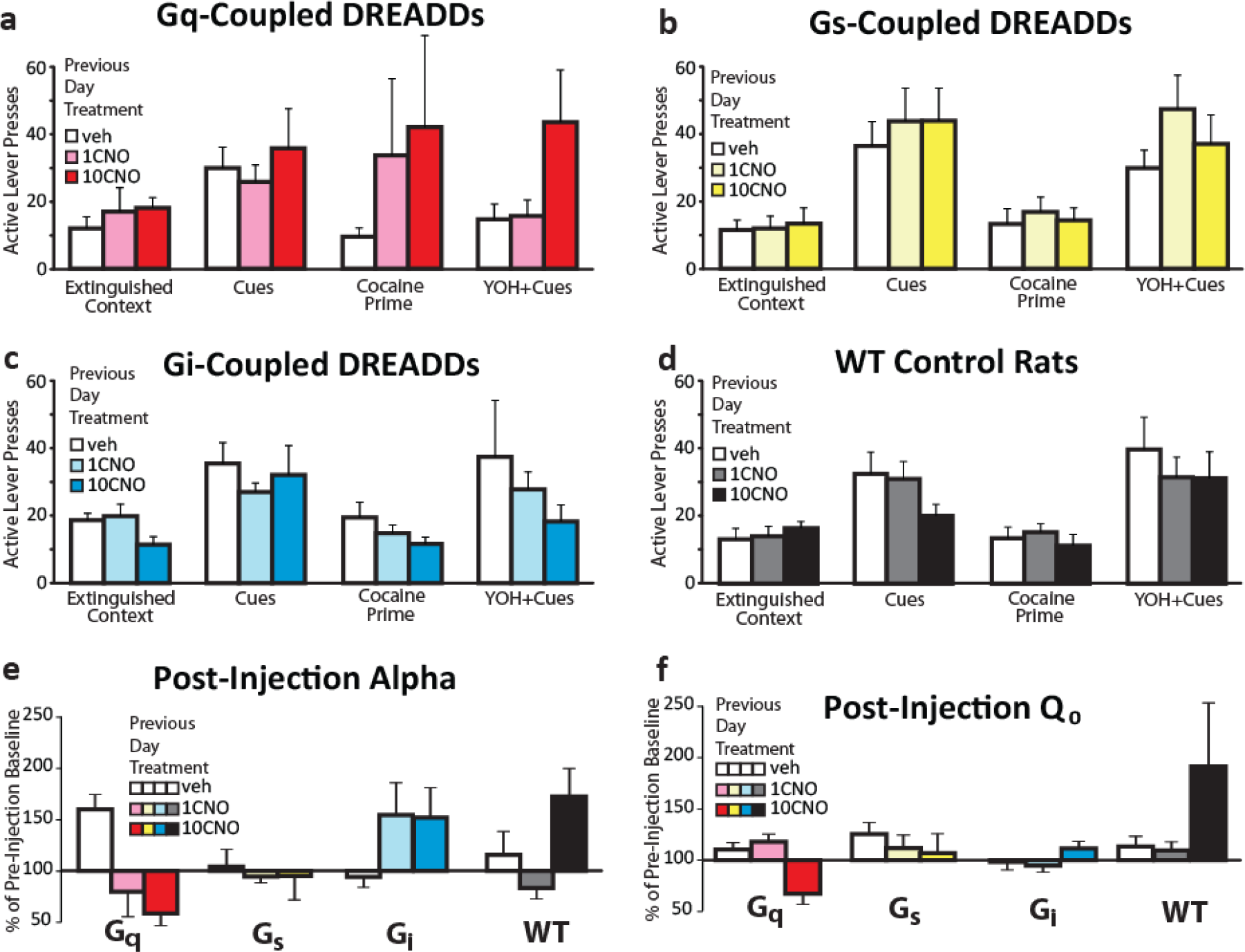
CNO Does Not Have Persistent Effects on Behavior. No consistent evidence for long-lasting effects of CNO in **a)** hM3Dq, **b)** rM3Ds, **c)** hM4Di, or **d)** WT control rats was found, as demonstrated with AL presses on the extinction re-training trials held 24hrs after injection of vehicle (white bars), 1mg/kg CNO (light red, yellow, blue, black bars), or 10mg/kg CNO (dark color bars; repeated measures ANOVA, p>0.05 for each test condition/DREADD group). Similarly, no persistent effects of prior-day CNO was observed on: **e)** demand elasticity (α) or f) free cocaine intake (Q_0_). Bars in panels **e-f** represent percentage of pre-injection baseline, on cocaine self-administration training days following tests days with veh or CNO (1 or 10mg/kg).

To characterize potential nonspecific motor effects of VTA DA neuron manipulations that could alter lever pressing, in a subset of each DREADD group we examined effects of systemic CNO on horizontal activity (Horiz.) and rearing in a pre-habituated test cage. CNO increased activity in hM3Dq rats across the entire 2h session (Horiz: F_1.042,5.21_=6.774, p=0.046; Rears: F_2,10_=6.038, p=0.019), trended toward decreasing activity in hM4Di rats (Horiz: F_2,16_=3.253, p=0.065; Rears: F_2,16_=2.239, p=0.139), and did not affect activity in rM3Ds or WT rats (rM3Ds Horiz: F_2,20_=1.43, p=0.263, Rears: F_2,20_=2.007, p=0.161; WT Horiz: F_2,12_=1.364, p=0.293, Rears: F_2,12_=3.129, p=0.081; **Fig.4e-f**). CNO locomotor stimulation persisted for at least 2h in hM3Dq rats (no interaction of Tx X Time: F_6,36_=1.184, p=0.337). Reinstatement effects of hM3Dq stimulation were stronger than locomotor activation effects in the same rats (mean cue reinstatement effect of 10CNO: 1116.4% of vehicle day, Horiz. locomotion effect: 361.5%), and the locomotion vs. motivation-enhancing effects of CNO in the same hM3Dq rats were not correlated (change from vehicle Horiz. not related to CNO effect on: cue reinstatement, r=-0.794, p=0.06; extinguished context, r=-0.494,p=0.32; prime: r=-0.539, p=0.27; YOH, r=-0.072, p=0.892).

## Discussion

Flere we systematically compared effects of chemogenetically manipulating VTA DA neurons, or their individual forebrain projections, during cocaine taking and seeking behaviors. These experiments revealed powerful, fundamentally modulatory roles for DA neurons in reinforcement, incentive motivation, and decision-making. Behavioral effects depended upon the GPCR signaling system and anatomical pathway that was experimentally engaged, as well as the behavioral and pharmacological context animals were in. Overall, results show that DA’s role in cocaine seeking is ultimately achieved by modulating behaviorally-relevant mesocorticolimbic circuits.

### Dual Roles for DA Neurons in High and Low Effort Cocaine Intake

Previous studies have indicated two, apparently opposing roles for DA in reward behavior: DA is involved in the subjective effects of psychostimulants, and blocking DA receptors causes rats to self-administer more cocaine to compensate(Suto and Wise, 2011; Willuhn et al., 2012). However, DA blockade also decreases effortful drug seeking in humans and animals leading to less cocaine intake (Berridge, 2007; Salamone and Correa, 2012). Here, we concurrently tested both of these putative roles for DA in a within-session cocaine behavioral economic paradigm, and show that DA neurons participate in both processes. At the start of each session, preferred cocaine levels are easily attained and cocaine intake during this period was increased when DA neurons were inhibited, or decreased when DA neurons were stimulated. This is consistent with the subjective effects of cocaine involving DA released from VTA terminals. However, later in the session rats were required to work harder to defend preferred cocaine blood levels. During this period, DREADD stimulation of VTA DA neurons increased responding, whereas inhibition in hM4Di rats showed less responding. This finding is consistent with the view that VTA DA drives motivation for cocaine. Therefore, these results indicate that VTA DA neurons are involved in both the subjective reinforcing, and motivational activating effects of cocaine, rather than playing a unitary role in reward. In plain language, when VTA DA neurons were stimulated rats wanted less cocaine but worked harder to obtain what they wanted, vs during inhibition of this DA system subjects wanted more cocaine but were less willing to work hard to obtain it.

### VTA DA Neuron Gq-Stimulated Cocaine Relapse

We find several distinct roles for VTA DA neurons in reinstatement behavior. Gq DREADD stimulation reinstated extinguished cocaine seeking in the absence of cocaine cues, priming injections, or pharmacological stress—indicating sufficiency of DA neuron Gq signaling for cocaine seeking. Gq augmentation of reinstatement behavior was strongest in the presence of response-contingent cocaine cues, suggesting a synergistic relationship between DA neuron stimulation and cue-induced cocaine seeking. Though DA stimulation also increased general locomotor activity, the magnitude of these effects was weaker than for stimulation of cocaine seeking, and was uncorrelated with motivation-enhancing effects in individual rats, indicating independent recruitment of these two distinct functions of VTA DA neurons.

Stimulating rM3Ds DREADDs in VTA DA neurons had only modest effects on cocaine seeking and other behaviors. Indeed, the only significant effect of rM3Ds stimulation was a mild increase in cue-induced cocaine seeking. In contrast, Gq stimulation in animals with equivalent DREADD expression dramatically increased cued cocaine seeking. As CNO enhanced DA neuron activity in rM3Ds rats (as measured by Fos and in vitro firing), and rM3Ds DREADDs have robust behavioral effects in other experiments(Ferguson et al., 2011; Ferguson et al., 2013; Gourley et al., 2016), our findings are unlikely to result simply from insufficient activity of the rM3Ds receptor. Instead, these findings reveal qualitatively different behavioral effects of Gq-versus Gs-signaling in VTA DA neurons, with preferential activation of motivation by Gq-stimulation. Although not assessed here, Gq- and Gs- activation could differentially modulate the firing patterns of DA cells or amount of DA released by them, as occurs with endogenous GPCR signaling in VTA neurons(Fiorillo and Williams, 2000; Foster et al., 2014). Also, given crucial roles for cAMP-dependent Gs signaling in DA neuron plasticity(Shen et al., 2013), and previous reports of Gs-DREADD stimulation causing persistent behavioral effects(Ferguson et al., 2013; Nakajima et al., 2016), it is also possible that Gs signaling plays a larger role in learning than in expression of previously learned motivation—this possibility should be tested in future studies.

### Chemogenetic Inhibition reveals behavior-specific roles for VTA DA neurons

Inhibiting DA neuron activity with hM4Di decreased some, but not all types of reinstatement behavior. hM4Di DREADDs reduced reinstatement elicited by either a cocaine prime or by cues plus the pharmacological stressor yohimbine. Given the robust modulation of conditioned reward seeking by VTA DA neurons, it is surprising that hM4Di VTA DA neuron inhibition failed to block cue-induced reinstatement. Instead, Gi inhibition altered the targeting of motivated behavior, increasing pressing of an inactive control lever (IL) that never yielded cocaine or cues. Increased responding on an inactive manipulandum for cocaine was also previously reported for systemic DA antagonist administration(Willuhn et al., 2012). hM4Di-induced IL pressing might therefore represent a shift to exploration of alternative strategies for attaining cocaine, but clearly does not demonstrate simple suppression of conditioned motivation. This intriguing finding points to a heretofore understudied role for VTA DA neurons in allocating effort to specific previously-rewarded actions during conditioned drug reinstatement, or in memory for cocaine associated-cues and their meaning.

### Functional Heterogeneity of DA Neuron Efferents.

VTA DA neurons show remarkable molecular and physiological diversity, related both to the efferent targets of individual neurons, and the behaviors these neurons regulate(Roeper, 2013; Lammel et al., 2014a; Barker et al., 2016). Here we demonstrate that stimulating DA projections to NAc, mPFC, or BLA is sufficient for enhancing the motivational impact of previously learned cocaine cues, but only DA terminals in NAc are sufficient to elicit reinstatement in the absence of discrete cues. We also found that stimulating DA neuron terminals in BLA enhances food intake, the first demonstration of BLA involvement in unconditioned feeding. We note, however, that chemogenetic stimulation of terminals of VTA TH(+) neurons via CNO microinjection also likely causes release of both DA and non-DA transmitters(Tritsch et al., 2012; Trudeau et al., 2014; Barker et al., 2016)—especially in mPFC, where DREADD expressing axons rarely co-expressed TH in these cocaine-experienced rats. Notably, intracranial CNO (1mM) effects are DREADD-specific(Mahler et al., 2014; Ge et al., 2017; Lichtenberg et al., 2017; McGlinchey and Aston-Jones, 2017) and are unlikely to result from non-specific effects of CNO or its metabolites at endogenous receptors(Gomez et al., 2017; Mahler and Aston-Jones, 2018). Together, these data show for the first time that presynaptic Gq stimulation of DA axons via CNO microinjection affects behavior, and does so in a pathway-specific manner.

### Effects of DREADD Manipulations on VTA DA Neuron Activity

Our extensive validation of bidirectional chemogenetic modulation of DA neurons in TH:Cre rats revealed several mechanisms through which activation of DREADD receptors modifies DA system activity. CNO increased activity of hM3Dq-expressing DA neurons as measured by in vitro firing and Fos expression, and also recruited otherwise quiescent populations of DA neurons in anesthetized rats, and indirectly reduced Fos in TH(-) VTA neurons. CNO also increased Fos and in vitro firing in rM3Ds rats, but did not affect Fos in TH(-) DA neurons, nor alter population activity in vivo. Finally, CNO in hM4Di rats failed to reduce firing or Fos in DA neurons under the measured circumstances, in which DA neuron firing was not robustly stimulated by endogenous behavior-related activity (in vitro slices, anesthetized rats, or homecage conditions for Fos).

We also examined the ability of DREADDs to presynaptically modulate NAc DA release, as is possible with inhibitory DREADDs in other neural circuits(Mahler et al., 2014; Stachniak et al., 2014; Buchta et al., 2017; Ge et al., 2017; Lichtenberg et al., 2017; McGlinchey and Aston-Jones, 2017). Electrically-evoked DA release was augmented by CNO in slices from hM3Dq rats, reduced in hM4Di rats, and unaffected in rats without DREADDs or with rM3Ds DREADDs, thereby validating the efficacy of axonally-applied CNO to presynaptically modulate DA release via Gi/o or Gq-coupled signaling.

### Translational Implications of Chemogenetic Circuit Interventions

A key advantage of DREADDs for intervening in neural circuits is the potential for translation of this or related approaches to the mental health clinic. The proximate cause of psychiatric disorders involves abnormal function of neural circuits, and a chemogenetic approach could therefore be useful in correcting neural activity with potentially fewer side effects than traditional psychotropic drugs or chronically implanted hardware. Such approaches will require extensive preclinical evaluation, but initial primate studies indicate safety and efficacy of DREADDs for modulating behavior and circuit activity(Eldridge et al., 2016; Grayson et al., 2016). Indeed, the largest limitation to pursing this approach in the clinic may be determining circuits that, if successfully modulated, would treat chronic psychiatric disorders in humans. The present results illustrate that even selective circuit manipulations like those of DA neurons here can have nuanced, context-specific effects on behavior.

## Author Contributions

**SVM:** Conceived and designed all experiments, conducted behavioral, in vivo electrophysiology, and IHC experiments, analyzed data, wrote paper.

**ZDB:** Conceived and conducted slice voltammetry experiments, analyzed these data, analyzed in vivo physiology data

**BMC:** Conducted behavioral experiments, analyzed these data

**BB:** Conceived and conducted behavioral economic analyses

**WB:** conducted slice electrophysiology experiments, analyzed these data

**ZAC:** Conducted in vivo electrophysiology experiments.

**EL:** Conducted Behavioral experiments and analyzed these data

**MR:** Analyzed in vivo electrophysiology data

**MS:** Conducted axon colocalization experiments, analyzed these data

**JM:** Conducted behavioral experiments and analyzed these data

**AR:** conducted slice electrophysiology experiments, analyzed these data

**RAE:** Conceived slice voltammetry experiments, analyzed these data

**GAJ:** Conceived and designed all experiments, wrote paper.

## Acknowledgements

Christina Ruiz, Matthew Harberg, Hannah Schoch, and Phong Do for assistance with behavioral testing and Fos analysis, David Moorman and Peter Kalivas for helpful comments, NIDA Drug Distribution Program and NIMH Drug Supply Program for providing CNO. Research funded by PHS grants K99/R00 DA035251, R01 DA006214, P50 DA015369, F31 DA036989, T32 DA007288, R01 DA033342A, F31 DA042505, R01 DA031900, MUSC Department of Laboratory Animals Research Facilities Grant, Irvine Center for Addiction Neuroscience (University of California, Irvine).

## Extended Data

**Extended Figure 3-1:**
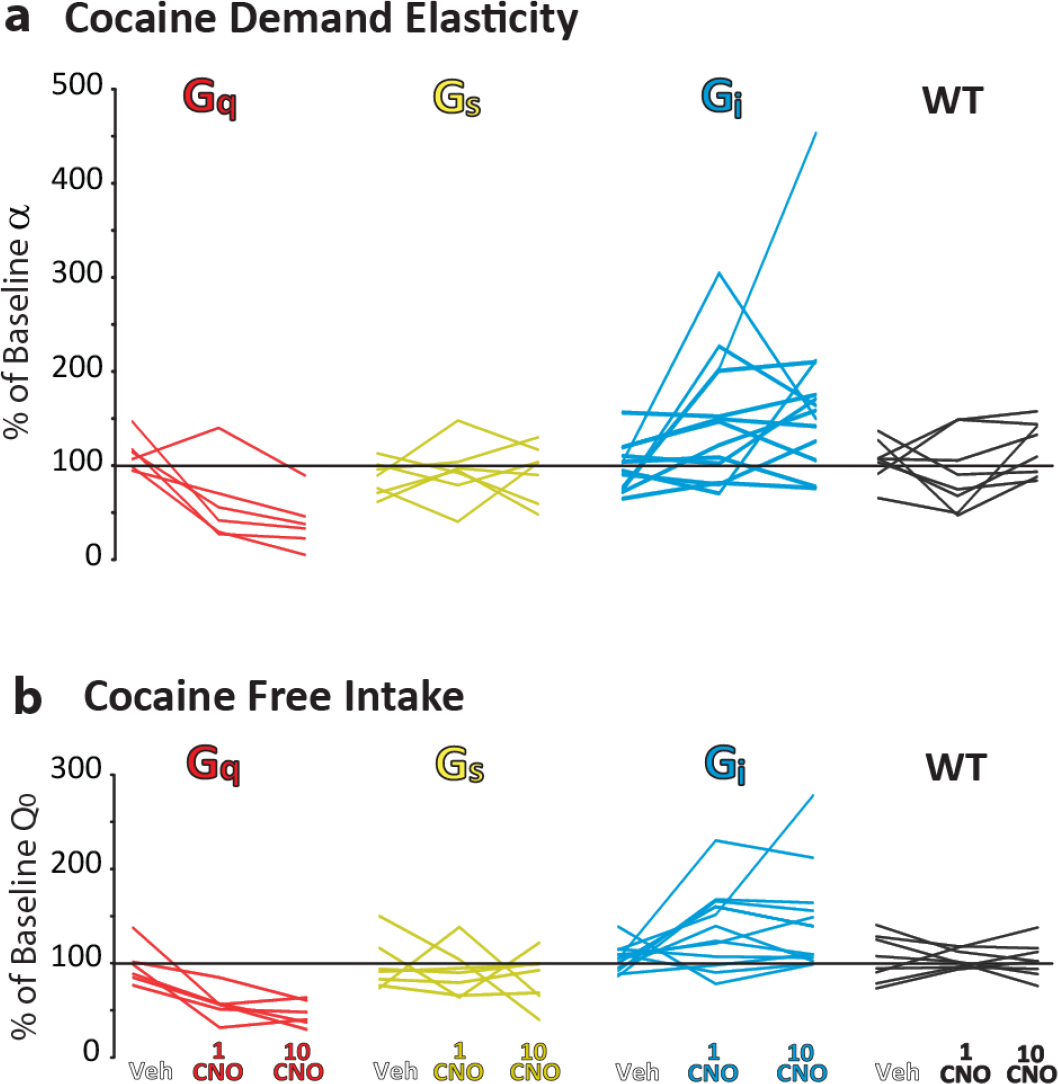
Rat-By-Rat Effects of Systemic CNO on Cocaine Demand and Free Intake. **a)** Cocaine demand elasticity (percentage change from stable baseline in α) after vehicle or CNO (1, 10mg/kg) is shown for each rat (1 rat=1 line). Leftmost point of each line represents vehicle day, middle=1mg/kg CNO, rightmost point= 10mg/kg CNO). Gq rats=red lines, Gs=yellow lines, Gi=blue lines, WT controls=black lines. **b)** Using the same symbol logic, cocaine intake at null effort (Q_0_; percentage change from stable baseline) is shown for each rat.

**Extended Figure 4-1:**
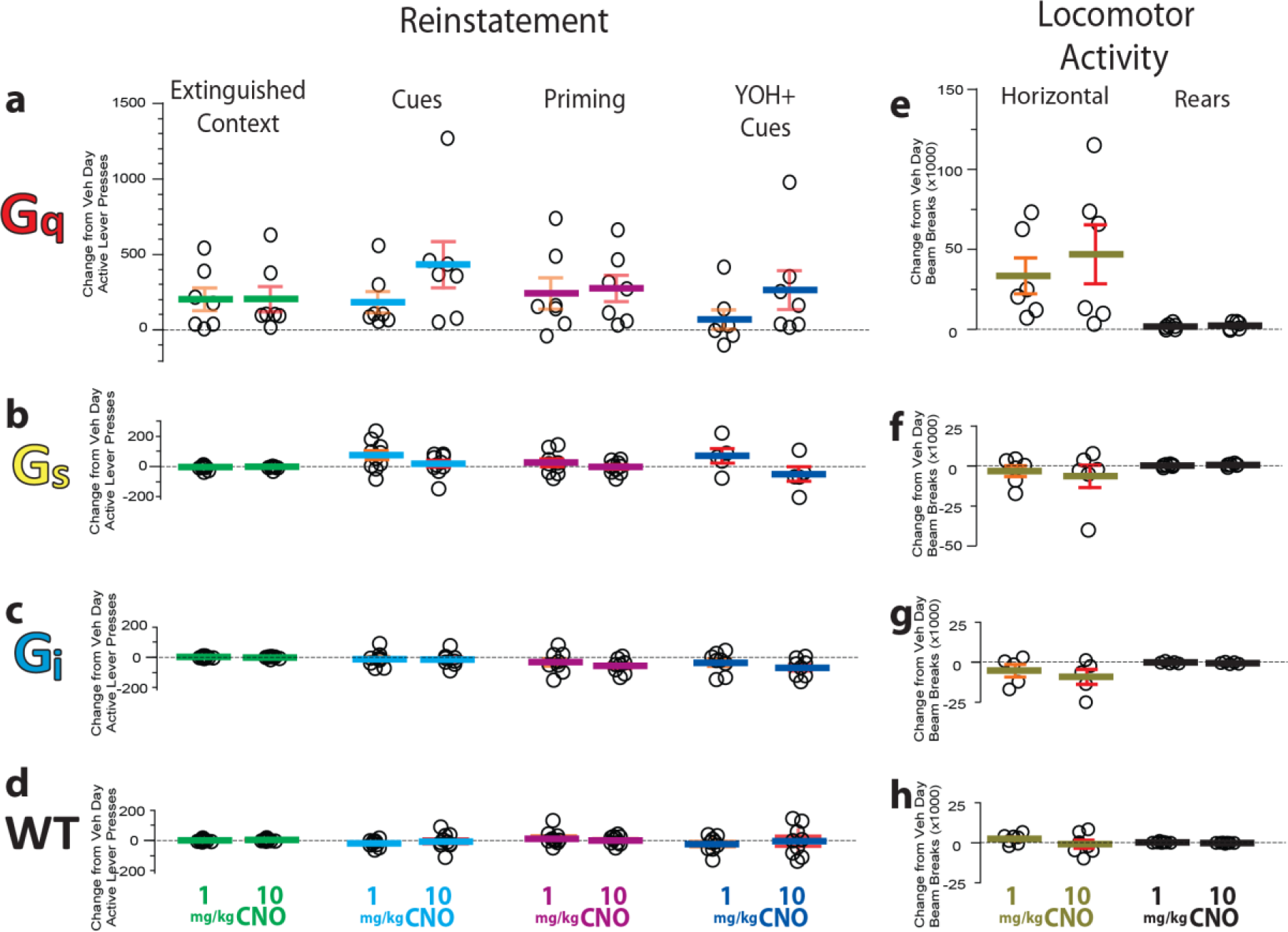
Reinstatement and Locomotor Activity Behaviors after Systemic CNO for Individual Subjects. Active Lever (AL) pressing is represented for each rat during reinstatement tests in each DREADD group: **a)** hM3Dq, **b)** rM3Ds, **c)** hM4Di, **d)** WT control. From left to right, change from vehicle day data (1 or 10mg/kg CNO) are shown for extinguished context, cue-induced reinstatement, cocaine primed reinstatement, and YOH+cue reinstatement. In panels **e-h**, horizontal locomotion (left) and rearing (right) is shown for 1 and 10mg/kg CNO days (change from vehicle day) for each rat. lines=mean, error bars=SEM.

**Extended Figure 7-1:**
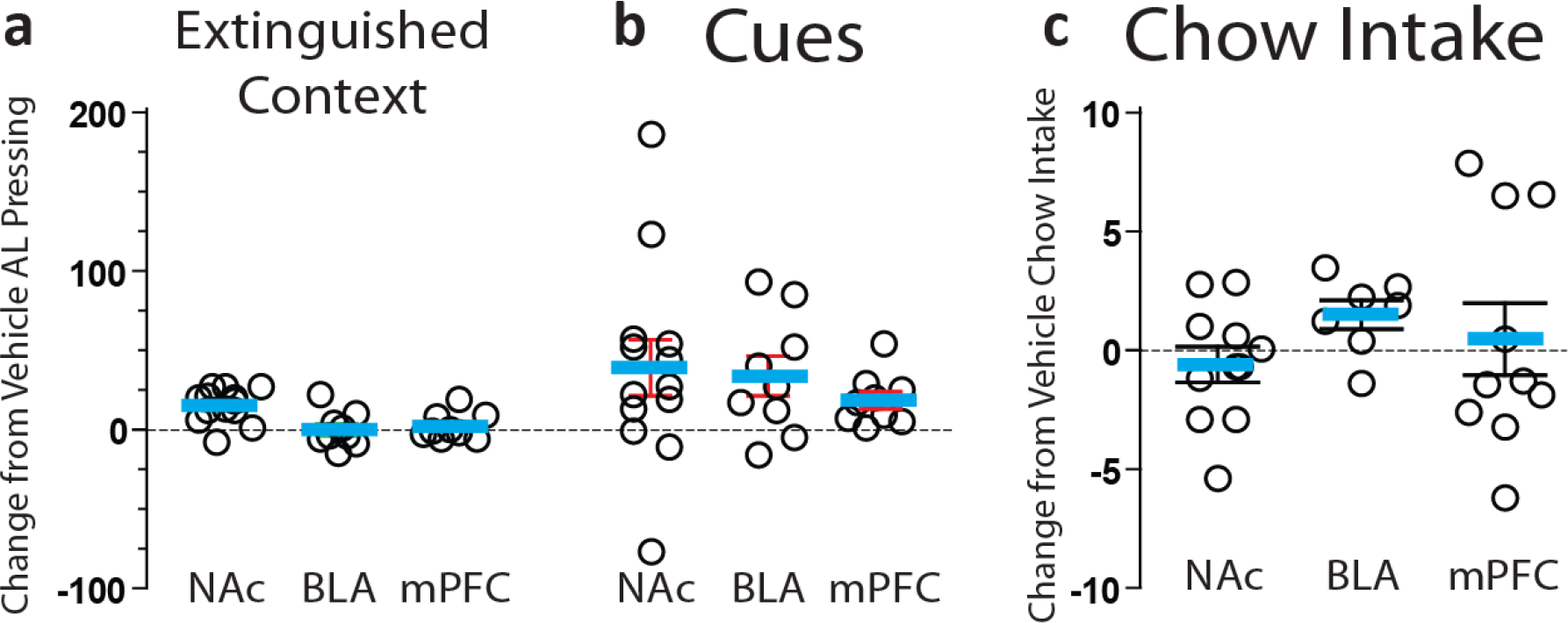
Pathway-Specific DREADD Stimulation Individual Subject Data. change from vehicle day AL pressing is shown for **a)** extinguished context and **b)** cue reinstatement. **c)** depicts change from vehicle day chow intake in a homecage free access test. In each panel, NAc, BLA, and mPFC means and SEMs are depicted, and individual rats are represented with circles.

